# Ultra-precise all-optical manipulation of neural circuits with multifunctional Cre-dependent transgenic mice

**DOI:** 10.1101/2021.10.05.463223

**Authors:** Hayley A. Bounds, Masato Sadahiro, William D. Hendricks, Marta Gajowa, Karthika Gopakumar, Daniel Quintana, Bosiljka Tasic, Tanya L. Daigle, Hongkui Zeng, Ian Antón Oldenburg, Hillel Adesnik

## Abstract

Determining which features of the neural code drive perception and behavior requires the ability to simultaneous read out and write in neural activity patterns with high precision across many neurons. All-optical systems that combine two photon (2p) calcium imaging and targeted 2p photostimulation enable the activation of specific, functionally defined groups of neurons in behaving animals. However, these techniques do not yet have the ability to reveal how the specific distribution of firing rates across a relevant neural population mediates neural computation and behavior. The key technical obstacle is the inability to transform single-cell calcium signals into accurate estimates of firing rate changes and then write in these cell-specific firing rate changes to each individual neuron in a targeted population. To overcome this challenge, we made two advances: first we introduce a new genetic line of mice for robust Cre-dependent co-expression of a high-performance calcium indicator and a potent soma-targeted microbial opsin. Second, using this line, we developed a pipeline that enables the read-out and write-in of precise population vectors of neural activity across a targeted group of neurons. The combination of the new multifunctional transgenic line and the photostimulation paradigm offer a powerful and convenient platform for investigating the neural codes of computation and behavior. It may prove particularly useful for probing causal features of the geometry of neural representations where the ability to directly control the topology of population activity is essential.

## Introduction

Many of the features of the neural code that underly sensation, cognition, and action remain unknown. Interventional tools that combine the simultaneous monitoring and manipulation of neural activity are proving essential to overcome this gap. Recently, all-optical techniques using 2p excitation have emerged as one class of tool for investigating the neural code with high spatial and temporal precision^1–9^. Under appropriate conditions, 2p optogenetics can achieve near single-cell resolution control over neural activity, thereby overcoming a major limitation of conventional optogenetic and electrical microstimulation^1–9^, helping to reveal basic rules of circuit connectivity and behavior^10–20^.

2p optogenetics as a technique, however, is not yet routine, and the lack of a simple, robust and convenient system to achieve widespread, stable co-expression of an optogenetic actuator (opsin) and an activity indicator in genetically defined neural populations has limited the use of all-optical techniques. Most 2p optogenetic studies have so far relied on viral expression^10, 11, 13, 14, 16–19, 21^ which typically requires invasive surgery, produces heterogeneous expression, can lead to toxic expression of the transgenes^22, 23^ or may directly impact synaptic physiology^24^. In contrast, transgenic lines provide convenient, stable and widespread expression of transgenes^25–28^, yet no transgenic line for the conditional co-expression of a calcium sensor and an optogenetic protein has been reported.

While experimenters have targeted specific, functionally defined neural ensembles, none have controlled the precise distribution of activity across the targeted neurons. Recent advances in the computational analysis of population activity have yielded new theories for how populations encode information and drive behavior. These include new insights into the geometry of neural representations and specific hypotheses about the importance of neural manifolds, communication subspaces, smoothness, and more^29–34^. Testing these new theories requires an approach that can enable perturbations that align with the estimated coding topology, deviate from it in defined ways, or control it outright. Doing so necessitates a system that can both read out and write in specific firing rates into individual neurons across a neural population. Existing all-optical techniques, however, have yet to achieve this for two reasons. First, it is not straightforward to infer the underlying spike rates measured with calcium sensors. Although a plethora of models exists for transforming calcium signals into estimates of the underlying spike trains, the biological variability of the spike-to-calcium signal transformation, even across neurons of the same cell type, constrains the performances of these models^35–40^. Second, it is challenging to ensure the optogenetic activation of specific numbers of spikes in targeted neurons due to a combination of the variability of opsin expression and the intrinsic excitability between neurons^16, 41^. Both of these issues imply that the models for inferring spike rates from calcium signals and for driving in specific spike rates with optogenetics must be precisely tuned for each neuron under study to achieve the desired neural manipulations.

To overcome these two obstacles, we developed a novel transgenic mouse line and a new paradigm for all-optical reproduction of precise activity patterns. For the genetic line we devised an allele that drives the co-expression of a high-performance calcium indicator, GCaMP7s^42^, fused directly to a potent soma-targeted microbial opsin, ChroME^41^. Sub-cellular targeting of the opsin to the somatic membrane is necessary to minimize off-target activation of unwanted neurons ^41, 43–46^. Because 2p stimulation requires higher energy than 1-photon (1p) stimulation, potent opsins, such as ChroME, are necessary to activate large ensembles of neurons without tissue damage ^21, 41, 47, 48^. Currently there are no transgenic lines that express ultrapotent opsins, and none that express a soma-targeted excitatory opsin. Among high potency opsins, ChroME has the fastest kinetics, which enables precise temporal control at high frequencies^41^, similar to its parent opsin, Chronos ^49, 50^. Among genetically encoded calcium sensors, GCaMP7s strikes a useful balance between the extreme sensitivity of GCaMP8s, and the much greater dynamic range and linearity of GCaMP6s^36, 42, 51^. We generated a single fusion protein between ChroME and GCaMP7s to ensure reliable co-expression of the two proteins and, via a single somatic targeting sequencing, simultaneously target both actuator and indicator to the soma. We leveraged the TIGRE2.0 system to achieve robust but non-toxic levels of expression of both transgenes^25^ and demonstrate that this mouse line provides high fidelity monitoring and optogenetic control of neural activity with 2p excitation.

Next, we combined the unique features of this mouse line with 2p holographic optogenetics to develop a new pipeline for precisely calibrating the transformation from 2p excitation pulses into spikes and spikes into calcium signals in individual neurons. We show that this multi-step, cell-specific calibration process enables the holographic recreation of precise sensory-driven population activity vectors by driving in firing rates tailored to each neuron in the targeted ensemble. Taken together, this new approach and this novel mouse line will empower neuroscientists to probe the precise geometry of neural codes and their relationship to behavior.

## Results

### Ai203 Transgenic Reporter Line

We first generated a novel transgenic reporter line using the TIGRE2.0 system which provides Cre- dependent co-expression of the potent, ultra-fast opsin, ChroME and the sensitive calcium indicator GCaMP7s ^25, 41, 42^. We used the Kv2.1 soma-targeting (st) sequence to enrich opsin expression at the soma, as described previously ^25, 41, 43, 44, 46, 52^. To ensure co-expression of the calcium sensor and the opsin, we fused GCaMP7s directly to the opsin (‘st-ChroME-GCaMP7s’), producing a soma-targeted calcium indicator ^53, 54^. The construct also includes a FLAG tag for antibody labeling in post-mortem tissue when necessary. We termed this line, TITL-st-ChroME-GCaMP7s-ICL-nls-mRuby3-IRES2-tTA2, ‘Ai203’ (Fig.1A and see supplementary note).

**Figure 1:**
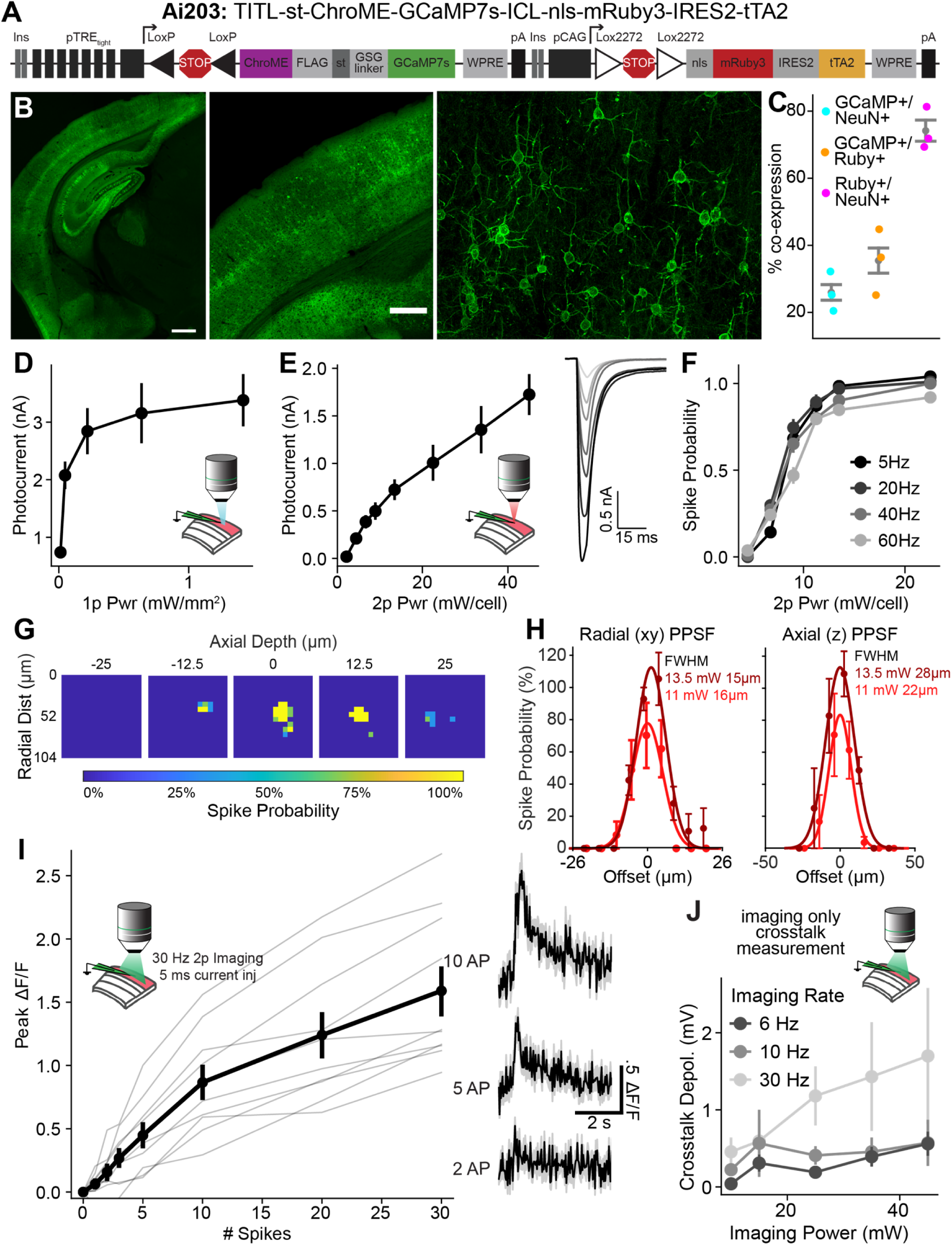
Characterization of expression, photocurrents, and GCaMP fluorescence in Vglut1-Cre;Ai203 mice. **A:** Schematic showing the Ai203 transgene. **B:** Confocal images of a representative Vglut1-Cre/wt;Ai203/wt heterozygous double transgenic mouse showing GCaMP7s (green) expression in primary visual cortex at different zoom. Images stitched from multiple fields of view. Far right, maximum z projection **C:** Quantification of co-expression of GCaMP7s, anti-NeuN and mRuby3 based on manual cell counting in visual cortex. n=3 mice. **D:** Photocurrents in opsin-positive L2/3 cells evoked by 1p illumination at 510 nm at varying powers. n=13 cells. **E:** Left, same as (A) but for 2p illumination. Right, averaged evoked photocurrent traces for an example cell, power increases from light to dark. n=11 cells **F:** Spike probability during 2p stimulation with a train of ten 5 ms pulses at different frequencies as a function of power. n=11 cells **G-H:** Assessment of physiological point-spread function (PPSF) of photostimulation-evoked spiking of opsin-positive neurons measured by loose-patch recordings. **G:** Heatmaps from an example cell depicting spike probability across five planes at power 25 mW. Depth z = 0 corresponds to the plane of the patched cell. **H:** Across cell mean PPSF for axial (left) and radial (right), at two powers. n=4 cells. **I:** Characterization of GCaMP fluorescence response. Simultaneous whole cell recording and 2p imaging was conducted and cells were electrically stimulated with trains of 5 ms current injections at 30 Hz. Peak ΔF/F fluorescence responses to different numbers of induced action potentials. Grey lines, individual cells. Black, average. n=10 cells. Right, traces for an example cell **J:** Mean depolarization induced by imaging laser when imaging at different frequencies with FOV size 980 x 980 µm and dwell time 46 ns/µm. n=4 cells. D-H,J: data are presented as mean ± sem. C,I: data are presented as mean ± bootstrapped 68% confidence interval

To test the new Ai203 transgenic line, we crossed it to the forebrain excitatory neuron driver, Slc17a7- IRES2-Cre (henceforth referred to as ‘Vglut1-Cre’) ^55^. Heterozygous double transgenic Vglut1-Cre;Ai203 mice exhibited high levels of st-ChroME-GCaMP7s expression across the entire forebrain (Fig. 1B, Fig. S1). As has been noted for some other combinations of Cre-driver and TIGRE2.0 reporter lines, the TRE-dependent st-ChroME-GCaMP7s was only expressed in a subset of excitatory neurons even though Vglut1- IRES2-Cre labels nearly all forebrain excitatory cells ^25, 55^. In contrast, the CAG-driven nls-mRuby3 was densely expressed, consistent with it labeling a large fraction of Cre+ glutamatergic cells (Fig. 1C, Fig. S1; supplementary note). Quantification of expression in primary visual cortex (V1) showed that 35 ± 6% of mRuby3+ cells were GCaMP7s+ (Fig. 1C, n=3 mice), implying that about one third of cortical excitatory neurons expressed the st-ChroME-GCaMP7s transgene.

We next examined expression when we crossed Ai203 to other Cre lines. Cux2-Cre-ERT2;Ai203 mice displayed robust expression across the upper layers of cortex, as expected for this Cre driver (Fig. S2A-D). GCaMP7s appeared to be largely restricted to the membrane of the soma and the proximal dendrites (Fig. S2E-F). To examine Ai203 expression in other neural populations, we crossed Ai203 mice with Vgat-IRES-Cre mice to label all GABAergic neurons ^56^. We observed brain-wide expression in inhibitory neurons. In the cortex, GCaMP7s expression appeared weaker in Vgat-Cre;Ai203 mice than in Vglut1-Cre;Ai203 mice but was strong in the striatum and cerebellum (Fig. S2G-J). Thus, Ai203 appears to express well across many neural populations, but, like many other TIGRE2.0 lines^25^, may not express strongly in cortical inhibitory neurons.

### Photostimulation, calcium imaging and circuit mapping in Ai203 mice *in vitro*

A key test of this new line is whether st-ChroME-GCaMP7s is expressed at sufficient levels for both photostimulation and calcium imaging under a variety of conditions. Under conventional 1p illumination, we observed strong photocurrents with peak amplitudes of 3.4 ± 0.4 nA when measured in voltage clamp at -70 mV (Fig. 1D, n=13 cells). Using 2p illumination (via 3D Scanless Holographic Optogenetics with Temporal focusing (3D-SHOT), see Pegard et al., 2017 and Mardinly et al., 2018 for details) peak photocurrents were 1.7 ± 0.2 nA, comparable to that previously reported for virally expressed ChroME (Fig. 1E; n=11 cells)^41^. To test if these 2p-induced photocurrents were sufficient to evoke spiking, we made cell-attached recordings and illuminated the cells with trains of ten 5 ms light pulses at varying frequencies.

Indeed, this reliably drove spiking at high frequencies (60 Hz) and at low powers (13.5 mW) (Fig. 1F; n=11 cells). Neurons followed pulse trains with sub-millisecond jitter (0.86 ± 0.04 ms; n=11 cells) across all stimulation frequencies tested. These data demonstrate that Ai203 mice provide robust expression of soma-targeted ChroME that drives temporally precise, high-fidelity spiking with 2p activation.

To test the effective spatial resolution of 2p holographic stimulation in Ai203 mice, we recorded light-evoked spiking from opsin-positive L2/3 pyramidal neurons in loose patch while stimulating each point in a small 3D grid surrounding the cell (Fig. 1G). We found an effective spatial resolution on par with previous reports using 3D-SHOT with soma-targeted opsins (Fig. 1H; FWHM @ 13.5 mW = 15 ± 2 μm radial; 28 ± 4 μm axial, n=4 cells) ^41, 57^.

Next, we asked whether the opsin-fused and soma-targeted GCaMP7s sensor is capable of sensitively reporting neuronal activity. We made whole-cell current clamp recordings from st-ChroME-GCaMP7s+ L2/3 neurons in brain slices, and induced spiking with short (5 ms) current injections through the patch pipette while measuring GCaMP7s fluorescence using 2p calcium imaging. We observed robust fluorescence responses for even small numbers of action potentials (Fig. 1I). Peak ΔF/F was 0.9 ± 0.1 for 10 action potentials (Fig. 1I, n=10 cells).

In all-optical read/write experiments, the scanning laser used for calcium imaging can cause unintended activation of the opsin, a phenomenon termed imaging “crosstalk” ^21, 41, 58^. To quantify crosstalk in Vglut1- Cre;Ai203 we made whole-cell current clamp recordings and recorded voltage changes induced by 2p imaging. Under conditions commonly used for volumetric imaging in all-optical experiments (FOV 980 × 980 μm, imaging rate 6-10 Hz, 46 ns/μm dwell time), mean depolarization caused by the imaging laser was <1 mV (Fig. 1J, Fig. S3, n=4 cells). These experiments show that the st-ChroME-GCaMP7s fusion is compatible with 2p laser scanning imaging and exhibits strong fluorescence responses with similar kinetics to cytosolic GCaMP7s during supratheshold neural activity.

Mapping synaptic connectivity is fundamental to obtaining a mechanistic understanding of neural circuits. 2p optogenetic stimulation enables high-throughput stimulation of individual cells to test for monosynaptic connections^14, 43, 45, 59–62^. Most prior studies relied on viral expression of the opsin which could lead to biases in the obtain connectivity maps due to incomplete spatial spread of the viral vector or due to direct physiological changes associated with AAV infection^24^. Thus, we tested the use of Ai203 for *in vitro* synaptic connectivity mapping using 2p photostimulation. To activate and identify putative presynaptic inputs, we stimulated individual spots in a larger 3D grid (Fig. S4A) while performing whole-cell voltage clamp recordings from an opsin-negative L2/3 pyramidal cell. We consistently identified between 11-15 putative monosynaptic connections to the patched cell within a single plane (Fig. S4B-C). These data demonstrate that 2p holographic optogenetics applied to the Vglut1-Cre;Ai203 transgenic line is suitable for mapping intracortical excitatory synaptic connectivity rapidly and volumetrically across brain tissue. Thus, Ai203 mice can allow investigators to probe synaptic connectivity without the potential pitfalls of viral expression.

### Visually evoked calcium responses and 2p optogenetic stimulation in Ai203 mice

To determine whether the expression of st-GCaMP7s in Ai203 mice is sufficient to report sensory evoked neural activity, we imaged calcium responses of L2/3 neurons in V1 through a cranial window (Fig. 2A) ^63, 64^. We observed robust st-GCaMP7s responses even at modest 2p imaging powers (∼50 mW at 920 nm). Cells expressing st-ChroME-GCaMP7s exhibited strong responses to high contrast drifting gratings (Fig. 2B). Importantly, a similar proportion of L2/3 cells in Ai203 mice were visually responsive compared to another transgenic mouse line which expresses cytoplasmic GCaMP6s (using Camk2a-tTA;tetO-GCaMP6s) and which were intracranially injected with an AAV encoding ChroME (percent visually responsive: Ai203, 39.6 ± 2.6%, n = 8 FOVs; Camk2a, 40.4 ± 2.4%, n = 12 FOVs, p=0.832, t-test, Fig. 2C). Furthermore, a similar fraction of cells were orientation tuned (percent orientation tuned: Ai203, 27.9 ± 1.3%, n = 5 FOVs; Camk2a, 33.3 ± 2.1%, n=8 FOVs, p=0.095, t-tests, Fig. 2D), and their orientation-selectivity was similar albeit slightly lower (OSI: Ai203, 0.56 ± 0.04, n = 8 FOVs; Camk2a, 0.66 ± 0.01, n = 5 FOVs, p=0.0167, t-test; Fig. 2E). Finally, we measured contrast response functions and found no difference between Vglut1- Cre;Ai203 and cytoplasmic GCaMP6s (p=0.682, 2-way ANOVA on contrast response curves, Fig. 2F). These data demonstrate that the Ai203 line is suitable for large scale calcium imaging of sensory evoked neural responses *in vivo*. They also indicate that the visual response properties of V1 neurons in Ai203 mice are largely normal, implying that the expression of the transgene does not appear to alter network properties.

**Figure 2:**
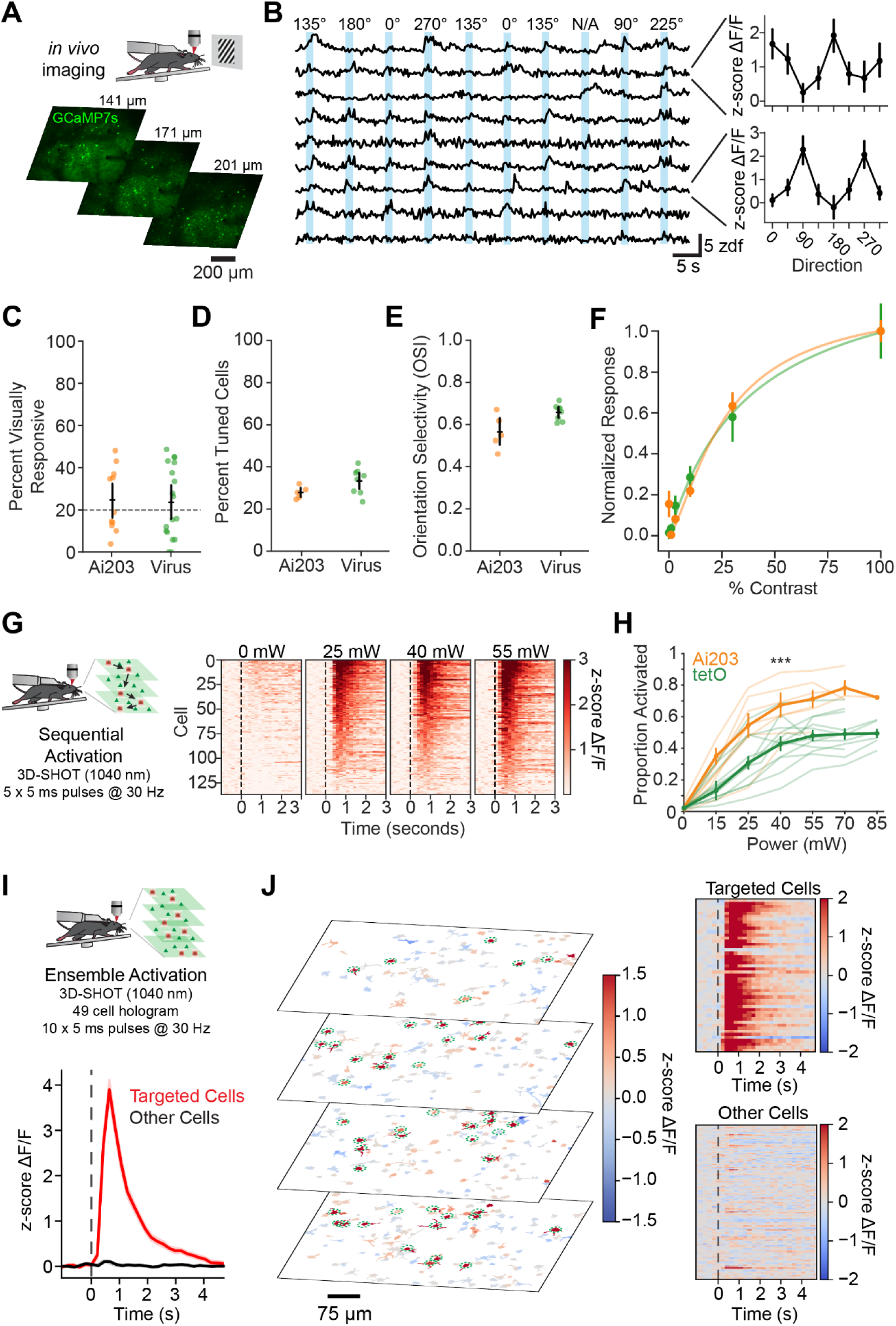
2p imaging of *in vivo* visually- and holographically-evoked responses in Vglut1-Cre;Ai203 mice. **A:** Top, schematic of in vivo imaging paradigm. Bottom, example fields of view for 3-plane (6.36 Hz volumetric) imaging. **B:** Left, example traces of fluorescence responses to drifting gratings in V1 pyramidal cells expressing st-ChroME-GCaMP7s. Right, direction tuning curves for 2 example cells. Scale bar applies to all example calcium traces. **C:** Comparison of percent visually responsive cells in Vglut1-Cre;Ai203 mice (orange, n=11 FOVs) and Camk2a-tTA;tetO-GCaMP6s mice expressing viral st-ChroME (green, n=18 FOVs). Mice expressing ChroME transgenically and virally had similar numbers of visually responsive, tuned cells (p= 0.854, t-test). FOVs with < 20% visually responsive cells were excluded from further analysis (dashed line). **D:** Comparison of percent significantly tuned cells in Vglut1-Cre;Ai203 mice (orange, n=5 FOVs) and Camk2a-tTA;tetO-GCaMP6s mice expressing viral st-ChroME (green, n=8 FOVs). Mice expressing ChroME transgenically and virally had similar numbers of visually responsive, tuned cells (p= 0.095, t-test). **E:** Comparison of mean orientation selectivity index (OSI) per session of significantly orientation-tuned cells, averaged by session for Vglut1- Cre;Ai203 mice (green) compared to Camk2a-tTA;tetO-GCaMP6s mice (p = 0.016). **F:** Contrast response functions (points) and corresponding Naka-Rushton fits (lines) to contrast modulated noise in Vglut-Cre;Ai203 mice (orange, n=6 FOVs) and Camk2a-tTA;tetO-GCaMP6s mice expressing viral st-ChroME (green, n=4 FOVs). Mice expressing ChroME transgenically and virally had similar contrast response functions (p=0.682, 2-way ANOVA). **G:** Left, schematic of single-cell holographic stimulation experiment. Targeted cells were sequentially activated across 3 z-planes. Right, heatmaps of z-scored ΔF/F responses at different powers for 110 significantly activated cells in an example session (16, 84, 82, and 110 cells were activated at 0, 25, 40, and 55 mW, respectively. one-tailed, rank sum test). **H:** Quantification of fraction of neurons activated at various stimulation powers during single-cell stimulation (p = 0.00012, repeated measures ANOVA). **I:** Schematic of ensemble stimulation experiment and average responses from targeted (left, red) and other non-targeted (black, right) cells. 49 neurons across 4 z-planes were activated simultaneously. **J:** Left, images showing individual cell responses during stimulation of 49 cells in a single hologram. Cell masks are colored by z-scored ΔF/F. Right, heatmap showing individual cells z-scored ΔF/F for targeted cells (49 cells, top) and all other cells (445 cells, bottom). 46/49 cells were significantly activated by stimulation (p<.05, one-tailed rank sum test). Data are presented as mean ± sem.

A primary use for the combined expression of st-ChroME and GCaMP7s is *in vivo* targeted optogenetic manipulation using multiphoton optogenetics. To first provide a basic validation for the utility of Ai203 in 2p optogenetics experiments *in vivo*, we used 3D-SHOT to holographically illuminate individual or ensembles of neurons while simultaneously imaging st-GCaMP7s responses using conventional multiplane 2p imaging. 2p photostimulation consistently drove large increases in st-GCaMP7s fluorescence in targeted cells (Fig. 2G-H, Video 1). Compared to mice intracranially injected with an AAV encoding st-ChroME (same mice as used above), Ai203 mice had a higher percentage of ChroME+ cells activated at all powers tested (proportion activated: Ai203, n=7; Virus, n=12, p=0.00012, 2-way repeated measures ANOVA, Fig. 2H). This high sensitivity enabled the co-activation of large ensembles of neurons with a single hologram, which we demonstrated by illuminating 49 L2/3 neurons simultaneously (Fig. 2I-J). Even during this large ensemble stimulation, neighboring non-targeted cells showed minimal activation, demonstrating that the Ai203 reporter is suitable for large-scale, high resolution photostimulation (Fig. 2J).

### Electrophysiological properties, and visual behavior in Ai203 mice

To examine the electrophysiological properties of cortical neurons in Ai203, we used whole-cell patch clamp recording in brain slices. We patched cortical pyramidal neurons in V1 of Vglut1-Cre; Ai203 mice blind to their GCaMP fluorescence to avoid any biases in cell selection. Cells were divided post-hoc into opsin+ and opsin- by the presence of ChroME photocurrent. We observed no differences in AP threshold (Fig. S5A; opsin+: -40 ± 1 mV, n=16 cells; opsin-: -39 ± .7 mV, n=14 cells; p=.38, t-test), although we did find that opsin+ cells exhibited a slightly more depolarized resting membrane potential (Fig. S5B; opsin+: - 74 ± 2 mV, n=16 cells; opsin-: -80 ± 1 mV, n=14 cells; p=.002, t-test). However, resting membrane potential was not correlated with photocurrent, implying that the expression of the opsin-GCaMP transgene *per se* did not explain the depolarized resting voltage (Fig. S5C, slope=-0.016, r^2^=0.0142, p=.66, F-test). We further characterized synaptic properties by analyzing miniature excitatory post-synaptic currents (mEPSCs).

Opsin+ cells exhibited a slightly higher rate and amplitude of mEPSCs (Fig. S5D-F; inter-event interval: opsin+: 0.27 ± .06 s, opsin-: 0.28 ± .04 s; p<.0001, ks test; amplitude: opsin+: 10.1 ± .3 pA, opsin-: 9 ± 1 pA; p<.0001, ks test, n=8 cells). These results could be explained if opsin+ neurons in Vglut1-Cre;Ai203 represent a specific subtype of excitatory neuron with slightly different physiological features. Alternatively, expression of the opsin-GCaMP transgene might alter synaptic properties. Since opsin+ neurons exhibit normal contrast sensitivity and orientation tuning *in vivo* (Fig. 2C-F), these different properties might only occur following brain slicing, or not impact the normal functional of these neurons in the intact circuit.

Transgenic opsin and GCaMP expression is particularly advantageous in behavioral training paradigms where long-term stability is essential. However, it is important that transgene expression does not itself impact learning or behavior. Thus we trained Vglut1-Cre;Ai203 mice on a visual contrast detection task to test both their ability to learn an operant behavior and to quantify their visual contrast sensitivity as a basic metric of vision (Fig. S6A). Similar detection tasks are known to be V1-dependent ^65–67^. We then compared contrast sensitivity, quantified as detection threshold, of Ai203 mice to C57BL6 wildtype (wt) mice and Camk2a-tTA;tetO-GCaMP6s mice trained on the same task. Vglut1-Cre;Ai203 mice performed as well as wildtypes and better than the Camk2a-tTA;tetO-GCaMP6s strains (Fig. S6B-C Detection threshold across genotypes: one-way ANOVA p=0.004 with post-hoc Tukey test: Ai203 vs wt p=0.3; Ai203 vs Camk2a p=0.004; Camk2a vs wt p=0.2; n=16 mice). Thus, Vglut1-Ai203 mice learn and perform this visual detection task comparable to wildtype mice, implying that transgene expression does not negatively alter learning or basic visual function.

Finally, we asked if optogenetic stimulation in Vglut1-Cre;Ai203 mice could alter behavioral performance on the visual contrast detection task. Indeed, photostimulation via a 470 nm optic fiber over V1 through a cranial window induced a dramatic increase in the false alarm rate and the hit rate for weak stimuli (Fig. S6D; Hit rate: no light 62 ± 4%, light 75 ± 3%, p=0.019; False alarm rate: no light 23 ± 3%, light 56 ± 5%, p=0.0016; paired t-test; n=4 mice). In contrast, wildtype animals showed no change (Fig. S6F; Hit rate: n light 57% ± 1%, light 58 ± 2% p=.5; False alarm rate: no light 20 ± 1%, light 24 ± 2%, p=.25; paired t-test; n=3 mice). Similar results were observed from activation of excitatory neurons in V1 in other studies ^65, 68^. Thus, Vglut1-Cre;Ai203 mice can be used for probing the impact of cell-type specific optogenetic manipulations on behavior. Taken together with the previous findings, these data demonstrate that Ai203 mice are suitable for a diverse set of neurobiological experiments.

### Programming precise population activity patterns with all-optical calibration

Neural codes depend not only on which neurons fire at a given moment, but also on the distribution of firing rates across a given neural population. This distribution constitutes the geometry of the neural code. Few if any studies have been able to write in specific patterns of activity across neurons in a population to causally probe how this geometry affects downstream activity or behavior. We reasoned that we could employ 3D- SHOT in combination with 2p imaging to achieve this goal, and that the Ai203 line would provide a particularly convenient platform for designing and validating such an approach owing to its strong, reliable expression of GCaMP7s and ChroME. We thus leveraged Ai203 mice to develop a new procedure for writing in precise numbers of spikes into different neurons simultaneously to recreate specific population activity vectors.

We considered two major challenges to reading and writing precise numbers of action potentials in neurons when use using all-optical techniques. First, to optogenetically write in a specific firing rate to each neuron in a targeted ensemble using holographic illumination pulses, we must ensure that each illumination pulse drives approximately one action potential. If we can achieve this, we can easily drive different spike rates into different neurons by varying the number and rate of illumination pulses delivered to each neuron.

However, the heterogeneity of both the intrinsic excitability and opsin expression across neuron makes it difficult to know, *a priori*, how much laser power to direct to each neuron to drive one spike per illumination pulse. The second challenge is that reading out a specific firing rate from a calcium signal is likewise complicated by heterogeneity across cells in the transfer function between action potentials and GCaMP fluorescent transients. Even if this transformation is approximately linear, its gain can vary widely across neurons even of the same cell type. Therefore, we designed a two-step calibration process fitting these transformations in each neuron using rapidly acquired all-optical experimental data.

The goal of the first step of this process, called ‘power calibration’, is to accurately identify the laser power needed to elicit one spike per illumination pulse for each neuron in a FOV. We generate a dose-response function that optically measures the fluorescence response of a neuron to a set of 2p illumination pulses (5 pulses, 5 ms duration) of increasing laser power. Laser powers that are too weak will only drive spiking on a some of the illumination pulses (Fig. 3A), but laser powers above the necessary power will generally not evoke additional spikes when using the fast ChroME opsin (Fig. 1F^41, 69^). Across a large population of neurons, we found that most neurons (1057/1730 cells, 8 FOVs, 5 mice;) exhibited a monotonic and saturating response curve that was well fit by a modified ‘hill function’ (Fig. 3B-C; mean R^2^ = 0.620±0.006). Saturation of the calcium response, at least for most cells, is likely not due to saturation of the opsin *per se*, since fluorescence response saturation was reached at considerably lower powers than photocurrent saturation (see Fig. 1E). Instead, it more likely represents that each of the 5 laser pulses drove exactly one action potential, implying that the saturation point indicates the minimum power necessary to achieve reliable spiking for each illumination pulse. Of the well-fit cells, the mean target power was 34.8 ± 0.4 mW, although there was a large range (Fig. 3D, 7 to 89 mW, N=1057 cells, 8 FOVs, 5 Mice), emphasizing the importance of this individual per cell calibration. Thus, this all-optical calibration allows us to empirically determine the illumination power needed in each neuron to drive an arbitrary firing rate.

**Figure 3:**
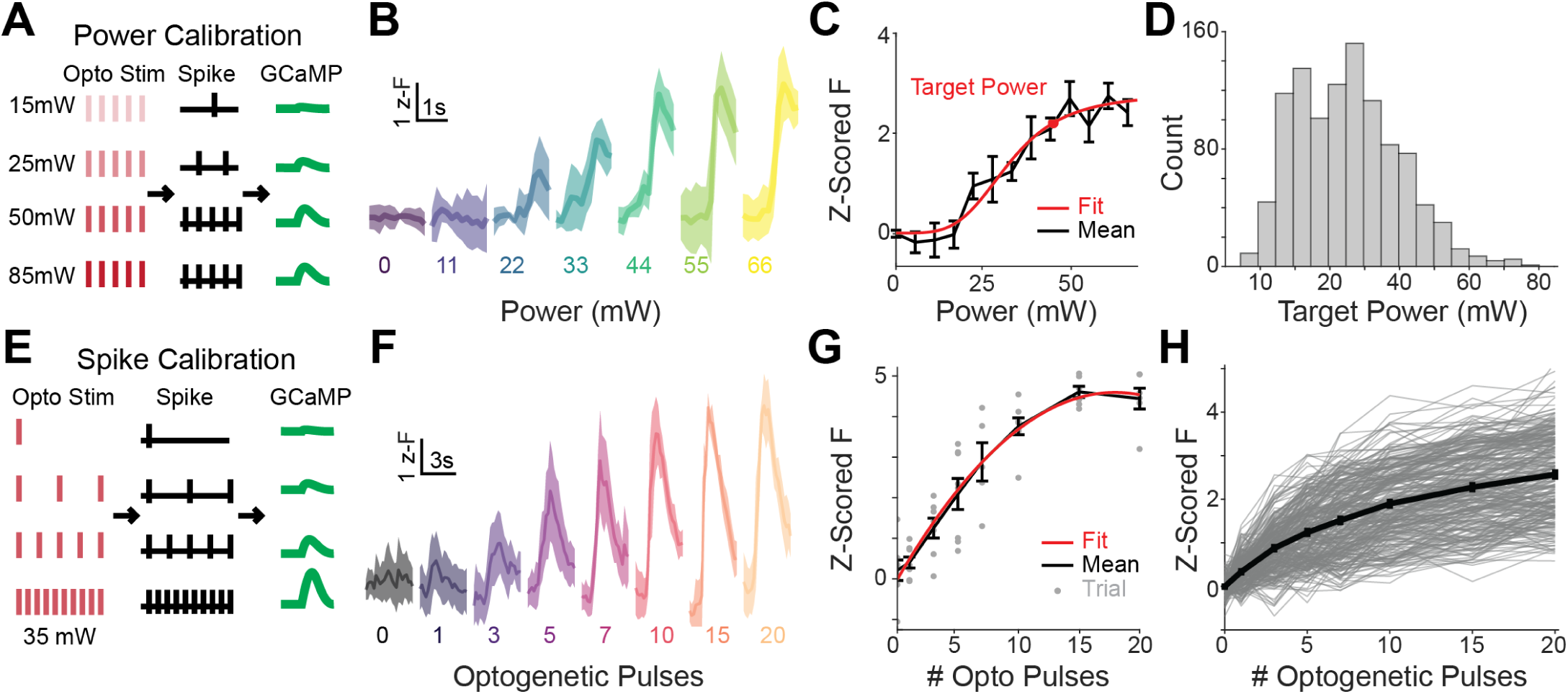
Calibration of spikes to GCaMP response. **A:** Schematic of the Power Calibration Procedure. **B:** Example cell stimulated with a train of 5 pulses at varying intensity. Mean ± 95% confidence Interval **C:** Mean response of an example cell, blue mean ± SEM. Fit line, red. With the calculated ‘target power’ noted in the red circle. **D:** Histogram of target powers of all fit cells, N=1057 cells, 8 FOVs, 5 Mice **E:** Schematic of the Spike Calibration Procedure. **F:** Example cell stimulated with trains of different number of pulses at the target power. Mean ± 95% confidence Interval. **G:** Mean response of an example cell, grey, mean ± SEM. Individual Trials marked in orange. Fit line, red. **F:** Overlay of 281 fit cells (light grey) Added Spike to DFF relationship (from 6 FOVs 4 Mice). Black, mean ± SEM of all fit cells.

The second step of this process, called ‘spike calibration’, maps the spike-to-calcium signal transformation in each neuron. In prior work, this transformation has been measured by using simultaneous electrophysiological measurement of action potentials or applied *post-hoc* with deconvolution algorithms^35, 36^. In order to estimate spiking on a per neuron basis within the same experiment, we employed an all optical, real-time approach to map spiking to calcium response across all neurons in the FOV. Using the power identified in the power calibration (above) we could use 2p optogenetics to write in known numbers of action potentials to each neuron. By varying the pulse and spike rate while simultaneously imaging the corresponding calcium transients we produced a response curve relating spike rate to fluorescence changes, which we fit with a quadratic function for each cell (Fig. 3E-G, see Methods).

Although cell-to-cell variability in the gain of each fit was high, as expected, 80% (281/350 cells) of targeted cells were well fit (Fig. 3H; R^2^ = 0.48 ± 0.01, N = 281 cells, 6 FOVs, 4 Mice).

We reasoned that these cell-specific calibration functions should allow us to recreate a natural population response by inferring and then holographically inducing a specific spike rate in each targeted neuron. To test this, we measured neural responses to drifting gratings at various orientations and selected a subset of the calibrated neurons that exhibited strong and orientation-tuned visual responses. Using our individually fit model of spikes to fluorescence, we calculated the number of optogenetic pulses necessary to recreate each neuron’s response to each grating, and generated a pattern of optogenetic stimulation that would drive the correct number of pulses in each neurons (Fig. 4A,D). We then used 3D-SHOT to drive in these computed light patterns. We observed a marked correspondence between the visually evoked activity and the holographically induced activity of the targeted neurons as intended (Fig. 4A-B). To quantify the success of this approach on a neuron-by-neuron basis, we computed the cosine similarity of the visual and the optogenetic response of each neuron. This analysis showed that the holographically recreated activity patterns were very similar to their natural counterparts (Fig. 4C; mean similarity: recreated: 0.61 ± 0.04; shuffled: 0.18 ± 0.04; p<3e-17 rank sum test; n = 155 cells, 5 FOVs, 3 mice).

**Figure 4:**
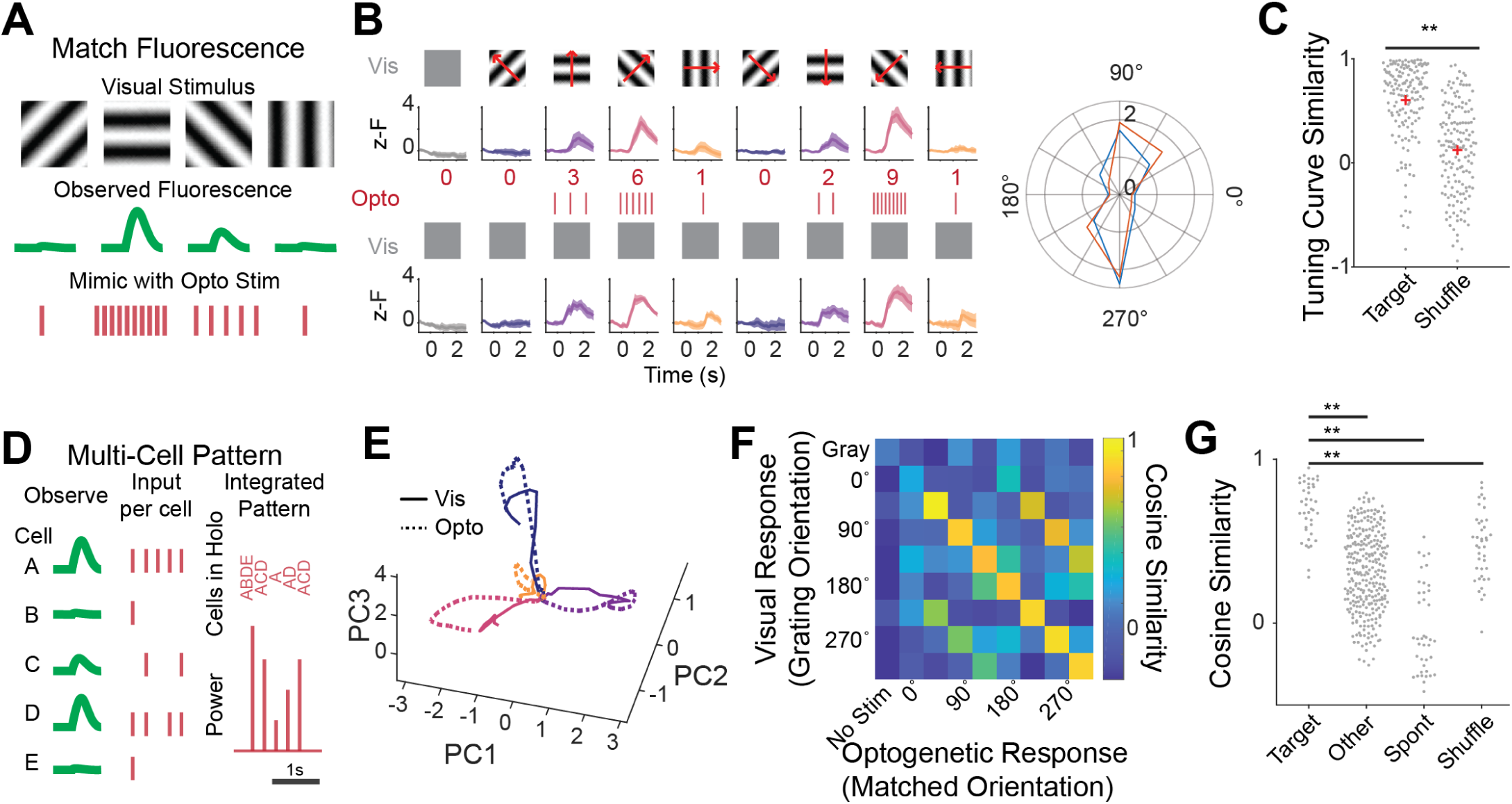
Reproducing naturalistic responses with 2p stimulation. **A:** Schematic of the decoding strategy. **B:** Example cell from a mouse presented with 8 drifting gratings or a grey screen and Mean ± 95% confidence interval of that cell’s response (Top). The calculated number of pulses to recreate that response in the context of a grey screen below, with the resulting response in the example cell (bellow). Right, polar plot of the cells’ Tuning curve visually evoked (blue), or by optogenetics alone (red). **C:** Tuning curve similarity of all visually evoked tuning curves vs the attempted optogenetically evoked tuning curves. Compared to cell shuffled controls. p<3e-17 rank sum test. N=155 cells, 5 FOVs, 3 Mice. **D:** Schematic of the Multi-Cell Pattern writing approach. **E:** Plot of population activity from all calibrated and used cells in an example field of view during visual stimulus (solid lines) or optogenetic only stimulation (dotted lines), for four drifting gratings, colors as in B. Dimensionality is reduced via principal component analysis. **F:** The Cosine Similarity between the visually evoked population activity and that evoked by optogenetic stimulation, for all tested holographic patterns from an example field of view. **G:** Cosine similarity of an ontogenetically evoked population vector to its target visually evoked pattern (‘target’), a different visually evoked pattern (‘other’), spontaneous activity (‘spont’), or a shuffled population (‘shuffle’). P values determined by rank sum test. Target vs other: p<5.5e-16, target vs spont p<5.5e-14, target vs shuffle p<1.9e-6. N= 40 optogenetically evoked patterns, 5 FOVs, 3 Mice. Ensembles contained between 19-43 cells.

At the population level, we saw appreciable similarity between the trajectories of visually induced and optogenetically induced activity patterns, when visualized with principal component analysis (Fig. 4E). To quantify their similarity, we computed the cosine similarity between the optogenetically evoked population vector and its visually evoked counterpart. Again, we found that optogenetic population vectors matched their intended targets well (cosine similarity 0.70 ± 0.03), much better than they matched other visual population vector from that FOV (0.32 ± 0.01, p<5.5e-16 rank sum test), spontaneous activity (-0.03 ± 0.04, p<5.5e-14 rank sum test), or a cell-shuffled dataset (Fig. 4F-G; 0.46 ± 0.03, p<1.9e-6 rank sum test. n=40 patterns, 5 FOVs, 3 Mice). These experiments establish and validate a general paradigm for recreating precise population vectors of neural activity in the intact brains of awake animals using all-optical read/write and 2p holographic optogenetics.

## Discussion

Here we report two advances: a novel multifunctional reporter line for all-optical circuit interrogation, and a new approach for recreating precise population activity patterns with 2p holographic optogenetics. The new mouse line provides the cell-type specific co-expression of an ultrapotent opsin, ChroME, and a high-performance calcium indicator, GCaMP7s, as a single soma-targeted fusion protein: st-ChroME-GCaMP7s. This is the first transgenic mouse that enables 2p all-optical interrogation of neural circuits with high spatial and temporal precision. Because the Ai203 transgenic line obviates the need for one or more viruses to co-express the opsin and the calcium indicator it will facilitate a variety of all-optical experiments by providing more reliable, reproducible, and stable transgene expression. This facilitated our development of a new paradigm that advances 2p optogenetic stimulation beyond the standard uniform activation of cells to reproducing naturalistic distributions of activity patterns in the population.

The robust, but stable levels of ChroME expression in Vglut1-Cre;Ai203 mice allowed us to photostimulate opsin-expressing neurons with even lower laser powers than mice expressing ChroME virally (via AAVs). This strong expression is helpful for a wide variety of experiments. *In vitro* we demonstrated use of the Ai203 line in 2p mapping experiments, which are a key method for understanding the link between the structure and function of neural circuits^14, 43, 45, 59–62^. *In vivo,* we demonstrated simultaneous activation of an arbitrary, experimenter-designated combination of nearly 50 neurons simultaneously *in vivo*, a number that exceeds what is sufficient to induce behavioral changes ^10, 13, 17, 19, 21^. Thus, the Ai203 line should be useful for probing causal relationships between ensemble activity and behavior. Additionally, the low powers required to reach reliable spiking allows greater flexibility in designing patterns of population stimulation, which is essential for our recreation of population activity.

Using the flexibility provided by the Ai203, we developed a new paradigm to artificially recreate population vectors of neural activity. In contrast to nearly all prior approaches which only activate specific cells or cell types without control over each stimulated neuron’s firing rate, or simply match fluorescence in closed-loop^70^, our new approach permits the user to distribute specific numbers of spikes to each of the target neurons arbitrarily or to recreate a measured stimulus-evoked response. Perhaps even more importantly, by using 2p holographic optogenetics to precisely calibrate the transformation from spikes into calcium signals in each neuron, the user can read out and write in firing rate distributions that approximate those driven naturally by sensory stimuli or during behavior. This new paradigm could be particularly useful for probing how the geometry of a neural code facilitates neural computations and drives behavior. A growing array of studies argue that neural coding operates along specific manifolds, through specific subspaces, or is organized into distinct topologies that might facilitate perceptual discrimination and generalization^29–34^. This new approach permits driving optogenetic perturbations that may align with specific aspects of estimated underlying coding topology, or deliberately deviate from it in specific directions to causally determine whether or how different dimensions of activity causally influence behavior.

Inferring underlying spiking from GCaMP fluorescence changes is a difficult problem due to the many sources of noise and variability within and across cells. Here, we sought to determine the spiking response of a neuron to a given stimulus so that we could reproduce an observed response. While we succeeded in matching optogenetically induced fluorescence to observed fluorescence in most cases, it is possible that our approach doesn’t necessarily match spike counts accurately. Various types of errors could lead to such inaccuracy: for example, high frequency spike bursts, which might generate larger calcium responses than low frequency firing of the same total spike count^35, 71, 72^, would lead to overestimates in the inferred spike rate, although prior work shows that most spikes in L2/3 pyramidal neurons, even in the awake state, do not occur in bursts^73^.

Spike inference models are another option for estimating the underlying spiking that give rise to a calcium response. Some approaches even try to estimate the underlying times on a single trial basis.

However, even state of the art methods suffer from many inaccuracies^37–40^ that may stem from cell-to-cell variabilities make estimating the calcium response to a single action potential challenging. The spike calibration approach we introduce here could be used in concert with deconvolution algorithms, especially for extending our approach to recreate precise spike timing. Currently, our system has limited ability to measure or recreate realistic spike timing since it is limited by the imaging frame rate and the kinetics of the calcium sensor. Incorporating faster imaging techniques^74^, faster calcium sensors^51^, or perhaps voltage sensors^75, 76^, as well as deconvolution models, can address this issue.

We note that while this process benefits from the Ai203 transgenic, it does not require it, and could be performed using standard viral approaches for transgene expression. However, the specific choice of opsin is likely to be crucial whether using transgenic or viral expression. Opsins with fast off-kinetics (such as ChroME) minimize the chance that single illumination pulses generate more than one spike because they allow the neuron to quickly repolarize, which is not true of opsins with much slower closing kinetics^41, 45, 77^. Shorter illumination pulse lengths may be able to overcome this for some slower opsins^21, 78^, but experimental validation of the power to spike transformation for any new experimental conditions would be required.

While Ai203 is a useful line for expressing ChroME, there are several important issues to consider when employing it for experiments. First, Ai203 provides only incomplete coverage of transgene expression in Cre+ cells. We found that when using the pan-excitatory driver Vglut1-IRES2-Cre ^55^, only about 35% of cortical excitatory neurons expressed the transgene. Other combinations of Cre driver and TIGRE1.0 and 2.0 reporter lines have shown this sparsity, which is thought to be due in part to promoter silencing of TRE (see supplementary note for further discussion) ^25, 27^. While sparse labeling precludes all-optical control and readout from many Cre+ neurons, it also offers advantages in many 2p optogenetic experiments (both in vivo and ex-vivo) due to reduced activation of neighboring cells ^1^. Incomplete expression is an issue with many transgenic expression strategies, such as TIGRE and thy1 lines, and new strategies to overcome this issue would greatly benefit the field.

A second consideration is the impact of the transgene on the physiological properties of the expressing cells, the network, and animal as a whole. *In vivo*, we found that st-ChroME-GCaMP7s^+^ cells in V1 exhibited normal responses to gratings of various contrasts and orientation tuning similar to another GCaMP transgenic line. The minor differences in orientation selectivity we did find could be due to differences in GCaMP type, levels or signal. At the behavioral level, Vglut1-Cre;Ai203 mice comparable contrast sensitivity to wild type controls, indicating that overall circuit function, at least in the visual system, is not compromised. In brain slices, we found that opsin+ cells exhibited slightly more depolarized resting membrane potential than opsin- cells, and also slightly larger and more frequent mEPSCs. This difference could be due to opsin+ cells representing a specific subtype of L2/3 pyramidal cell that differs from the unlabeled pool. Alternatively, these differences could be due to expression of the opsin or the GCaMP molecule *per se* changing physiological properties, although opsin levels were not correlated with change in resting membrane potentials. Regardless of the specific explanation, these putative differences appear to minimally impact circuit function *in vivo*.

The soma-targeted GCaMP construct here was produced by fusing GCaMP7s with an opsin that is soma-targeted via the Kv2.1 sequence. We observed robust fluorescence changes with GCaMP7s in response to small numbers of action potentials and reliably observed sensory-evoked activity *in vivo*.

Previous work with soma-targeted GCaMPs have shown that they reduce neuropil contamination, a common issue with both 1p and 2p calcium imaging^53, 54^. The soma targeting of GCaMP7s in the Ai203 line should likewise reduce contamination by signals from the neuropil. However, Ai203 mice should not be used with conventional 1p imaging since the illumination light for GCaMP7s will activate the ChroME opsin, causing substantial depolarization. Instead, Ai203 mice are well-suited for use with 2p laser scanning microscopy, and we found minimal optical crosstalk of the opsin by the 2p scanning laser under conventional imaging conditions.

The Ai203 reporter line and the ability to recreate population vectors both represent significant advances in all-optical experimental techniques. The more reliable and stable expression of the opsin and the calcium sensor in the Ai203 line compared to viral approaches will make all-optical experiments easier and more reproducible between animals and between studies. Recreating population vectors opens up a new range of experiments testing the geometry of neural codes, not just identity codes. Whether used together or separately, these advances are sure to promote new discoveries about neural coding.

## Acknowledgements

We thank the members of the Adesnik lab for comments, Savitha Sridharan for advice on construct development, Janine Beyer for technical support, and Nikhil Bhatla for assistance with histology. We thank Viviana Gradinaru and Ken Chan for the initial PhP.eB ChroME virus. Hillel Adesnik was a New York Stem Cell Robertson Investigator. This work was supported by the New York Stem Cell Foundation, National Institutes of Health (NIH) Grant UF1- NS107574 (H.A.), NIH Grant U19- NS107613 (H.A.), NIH Grant RF1- RF1MH120680 (H.A.), NIH Grant RO1- EY023756 (H.A.), National Science Foundation Graduate Research Fellowship (DGE 1752814 (H.A.B.)), NIH Grant F31-EY031977 (W.D.H.), NIH Grant K99-EY029758 (I.A.O.), NIH Grant U19-MH114830 (H.Z.). The content is solely the responsibility of the authors and does not necessarily represent the official views of the National Institutes of Health or the National Science Foundation.

---

**Supp. Figure 1:**
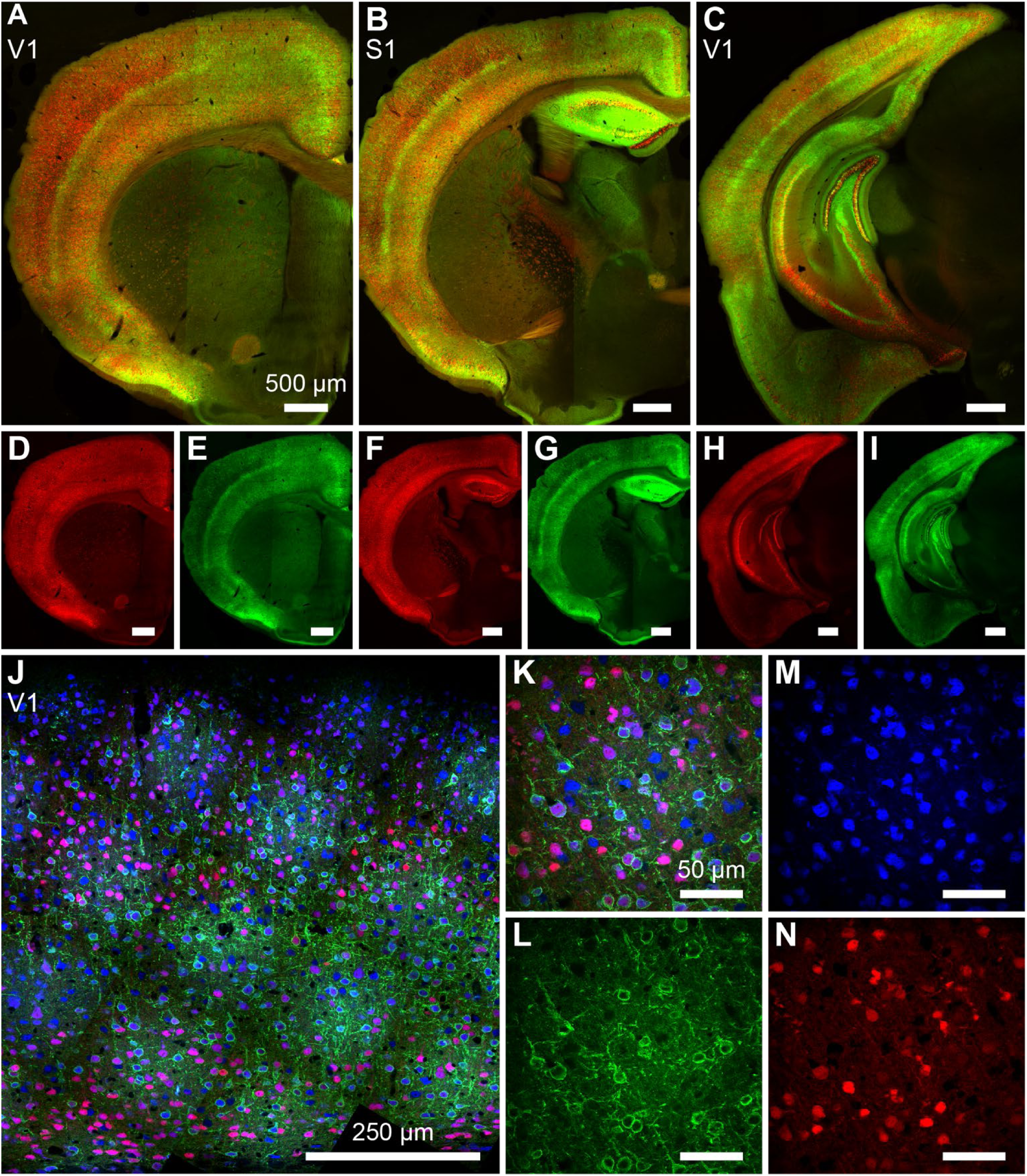
Expression of nls-mRuby3 and GCaMP7s in Vglut1-Cre;Ai203. **A-I:** Confocal images of brain sections from Vglut1-Cre;Ai203 mice showing expression of st-ChroME-GCaMP7s (green) and nls-mRuby3 (red) (images stitched from multiple fields of view). A-C, composite images of both mRuby3 and st-ChroME-GCaMP7s. D-I, sections in A-C separated by channel **J:** One of three areas used for quantification of expressing cells, showing in anti-NeuN (blue), nls-mRuby3 (red), st-ChroME-GCaMP7s (green). Area is composed of stitched fields of view taken at high magnification for good optical sectioning, then rotated and cropped to get a full depth section. **K-N**: one example field of view from the area in J. K: composite, M: anti-NeuN, L: st-ChroME-GCaMP7s, N: nls-mRuby3

**Supp. Figure 2:**
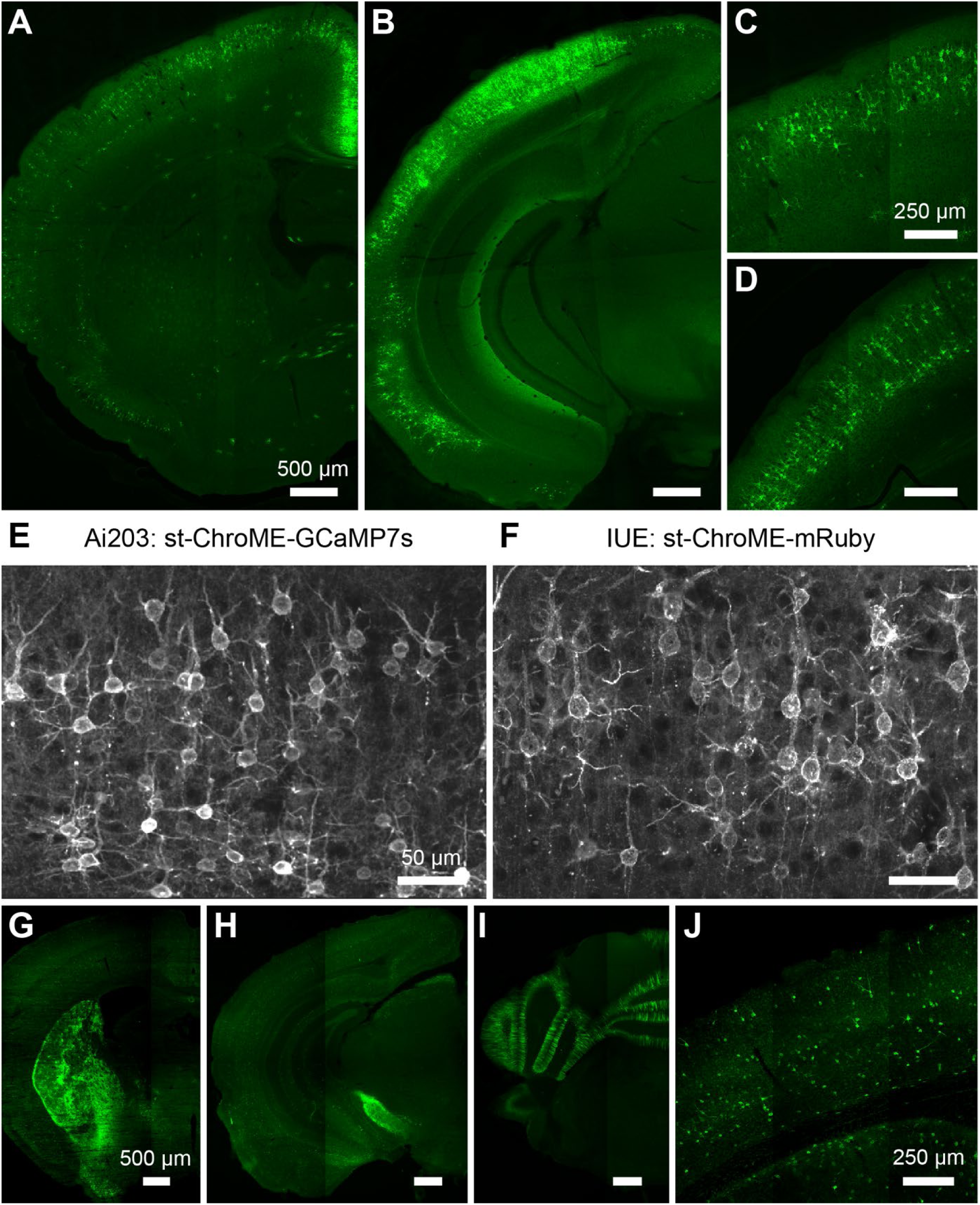
Expression of GCaMP7s and nls-mRuby3 in Cux2-Cre-ERT2; Ai203 and Vgat-IRES2-Cre; Ai203 mice. **A-D:** Confocal images of post-mortem tissue from one example Cux2-Cre-ERT2; Ai203 mouse at two different coronal sections and two zooms showing st-ChroME-GCaMP7s expression in green (images stitched from multiple fields of view). **E-F:** Comparison of in utero electroporation (IUE) st-ChroME-mRuby2 to transgenically expressed st-ChroME-GCaMP7s in Cux2-Cre-ERT2; Ai203. **G-J:** Confocal images of post-mortem tissue from one example mouse at different coronal sections showing st-ChroME-GCaMP7s expression in green (images stitched from multiple fields of view). **I:** Expression of st-ChroME-GCaMP7s in V1.

**Supp. Figure 3:**
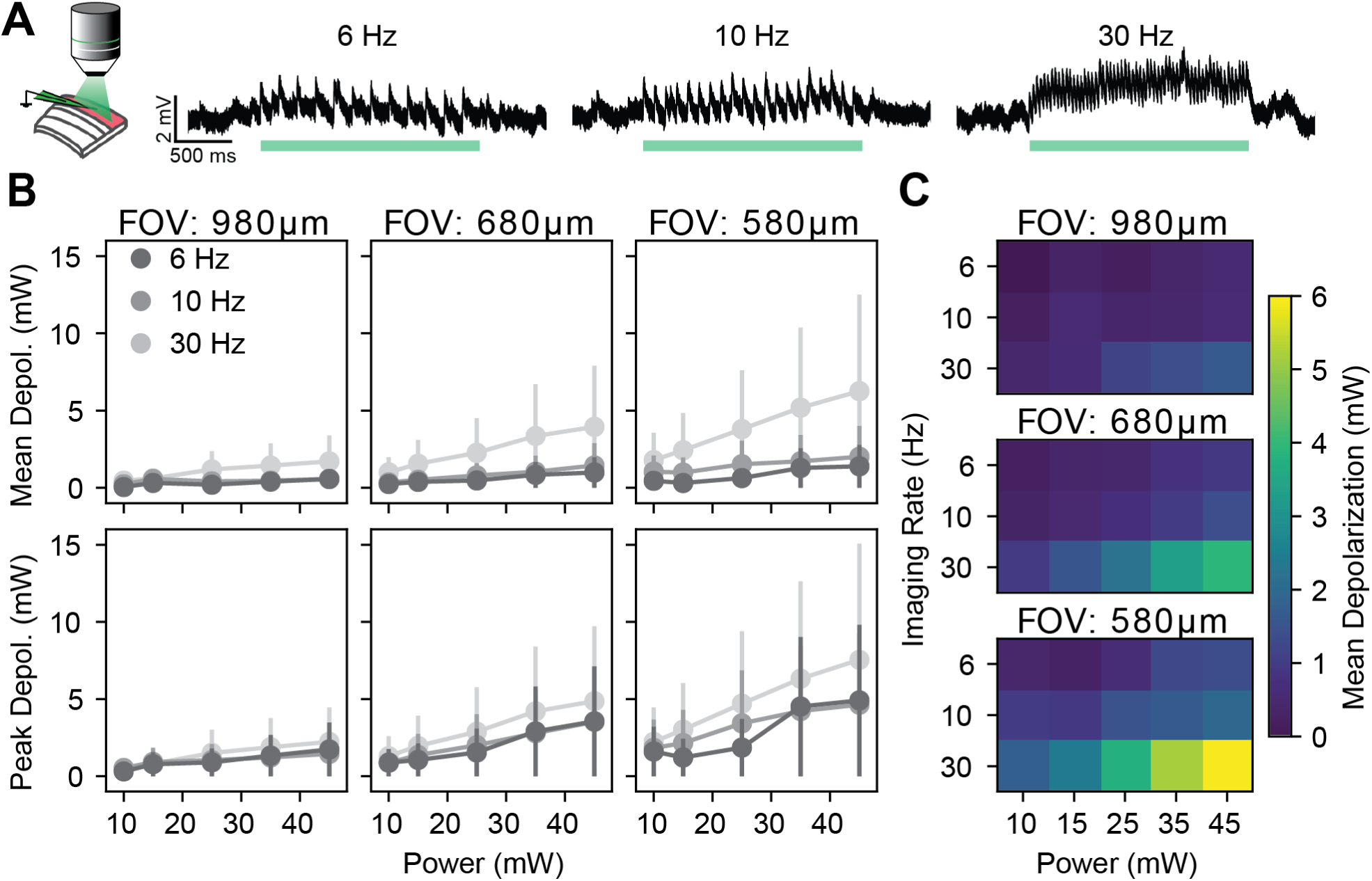
2p scanning-evoked depolarization *in vitro*. **A:** Left schematic of experimental setup. Whole-cell recordings were made during 2p imaging with different imaging rates, powers, and scanning field of view (FOV) sizes. Scanning-induced depolarization, or “cross talk,” was recorded. Right, 3 example traces from one cell recorded at different frequencies, all with 45 mW imaging power and a 980 μm FOV. **B:** Mean (top) and peak (bottom) depolarization during 2p imaging of opsin-positive neurons. From left to right, increasing zoom and decreasing field of view size, and increasing dwell times (FOV 980: 46 ns/um; FOV 680: 66 ns/um; FOV 580: 78 ns/um). Data are shown as mean ± sem. **C :** Same as in top of B, visualized as heatmaps.

**Supp. Figure 4:**
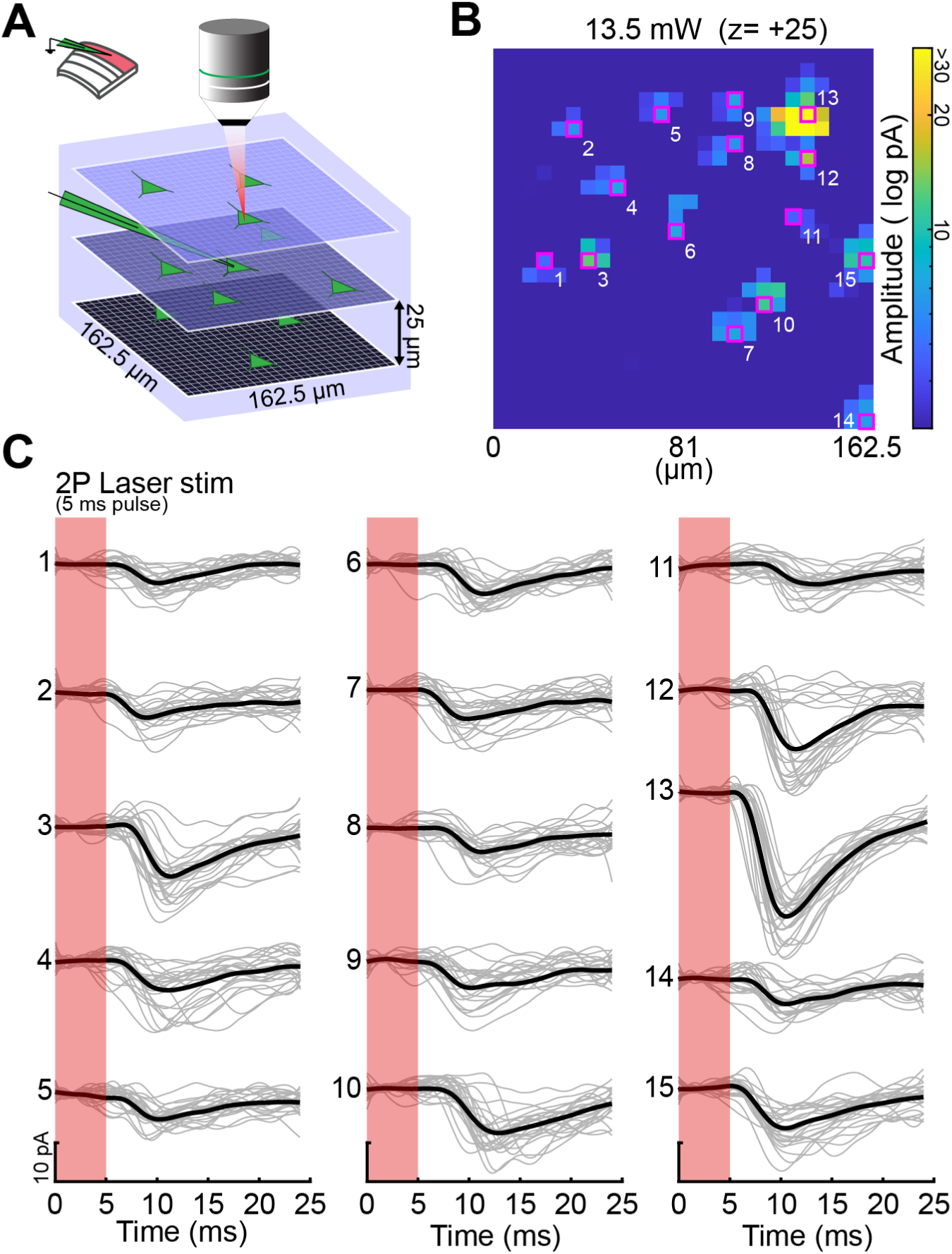
High resolution 2p holographic synaptic connectivity mapping in Ai203 mice. **A)** Schematic of protocol for “all-to-one” mapping of synaptic connectivity. 2p holograms were rapidly projected onto a 3D grid (162.5 × 162.5 × 100 μm) at 6.5 × 6.5 × 25 μm spacing (3380 total voxels) while an opsin-negative cell in the middle of the grid (*z* = 0) was voltage clamped to measure postsynaptic excitatory synaptic currents. **B)** Example heatmap of evoked postsynaptic EPSC amplitude by xy location of photostimulation sites in a plane z = 25 µm above the patched cell. The pixels with the highest average EPSC amplitude (across 20 trials) in 15 pixel clusters are highlighted in magenta. **C)** The photostimulation evoked postsynaptic responses (gray) and their average (black) from the 15 pixels selected in E. Data are presented as mean ± sem.

**Supp. Figure 5:**
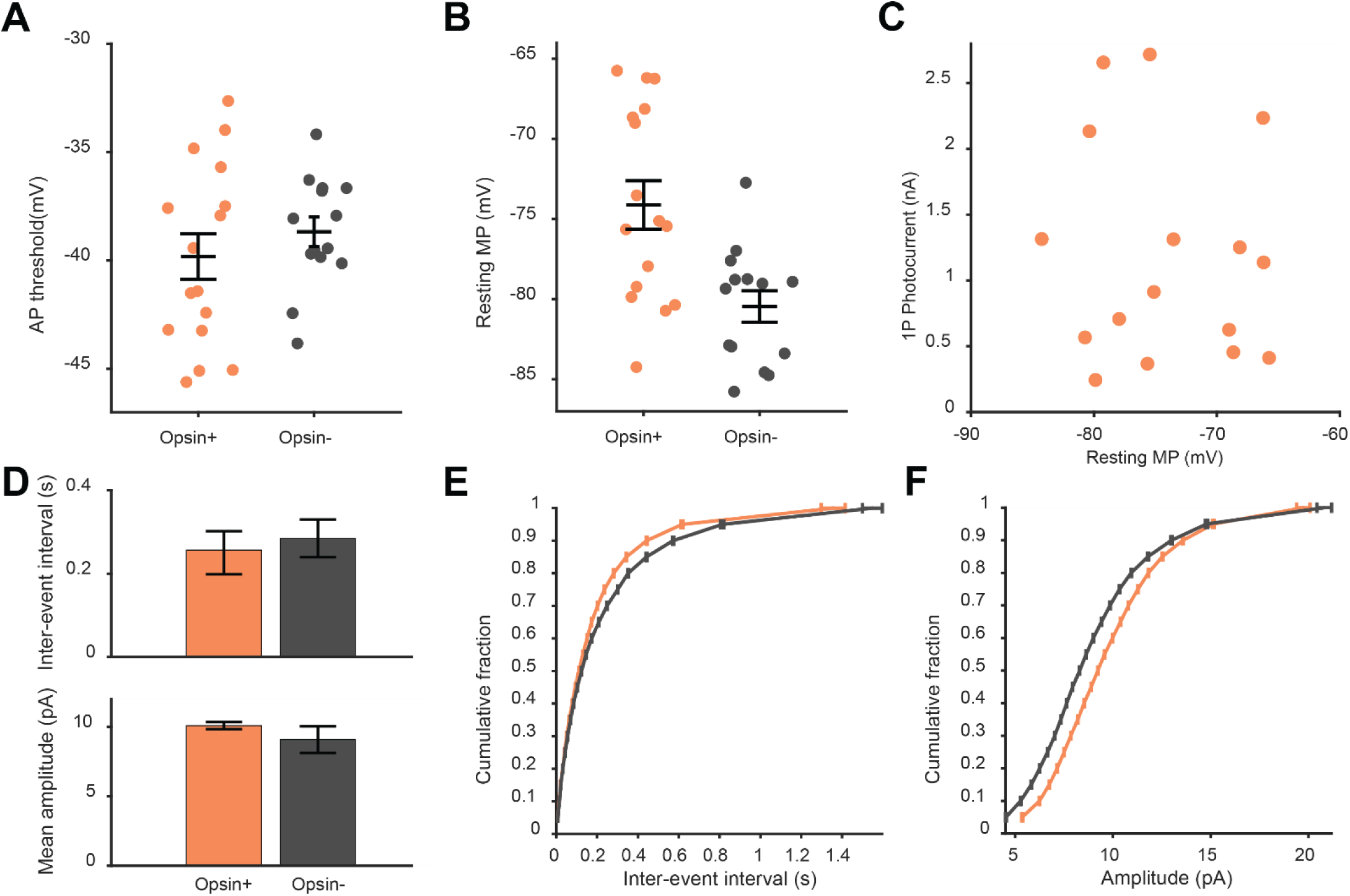
Comparison of basic physiological and synaptic properties of opsin+ and opsin- neurons in Ai203 mice. **A:** Action potential (AP) threshold f for opsin+ and opsin- neurons, categorized based on 1p photocurrent (p=.533; n=16 cells for opsin+ and n=14 for opsin-; t-test). **B:** Resting membrane potential (MP) for the cells in A (p=.002, n=16 cells for opsin+ and n=14 for opsin-; t-test) **C:** Magnitude of 1p photocurrent vs resting membrane potential for cells in A (slope=-.016, R^2^ 0.014, p=.66 that slope is different than 0) **D:** Comparison of miniature excitatory post-synaptic currents (mEPSCs) in opsin positive and opsin negative cells. Top, inter-event intervals between mEPSCS, bottom, mean amplitude of mEPSCs. **E:** Cumulative distribution of interevent intervals between opsin+ and opsin- cells. opsin+: 0.27 ± .06 s, opsin-: 0.28 ± .04 s; p<.0001, ks test. n=8 cells **F:** cumulative distribution of mEPSC amplitude between opsin + and opsin – cells. opsin+: 10.1 ± .3 pA, opsin-: 9 ± 1 pA; p<.0001, ks test, n=8 cells.

**Supp. Figure 6:**
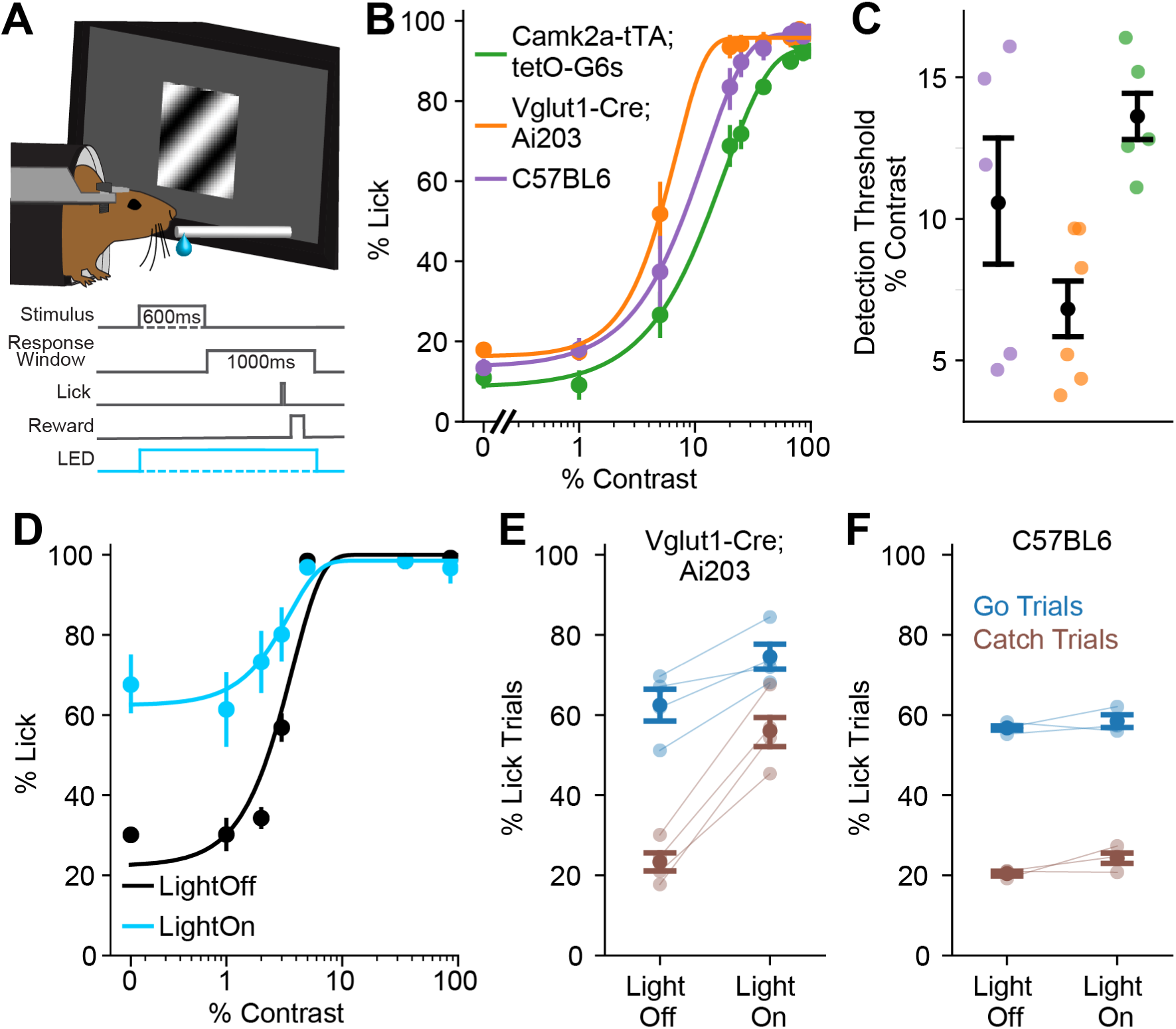
Optogenetic manipulation of operant behavior Vglut1-Cre;Ai203 mice alters performance. **A:** Schematic of the visual detection task. Gratings of varying contrasts were presented to mice trained to lick during a 1 second response window following stimulus presentation to indicate detection. **B:** Psychometric curves averaged across mice for of the three strains tested. n=6 mice for Vglut1-Cre;Ai203, n=5 each for wildtype and Camk2a-tTA;teto-GCaMP6s. **C:** Comparison of % correct, points are individual mice. % correct across genotypes: one-way ANOVA p=0.0025 with post-hoc Tukey test: Ai203 vs wildtype p=0.90; Ai203 vs tetO p=0.0050; tetO vs wildtype p=0.0056. **D:** Psychometric curves averaged across four sessions from one example mouse showing behavioral change with 1p stimulation at 470 nm of V1 excitatory cells. **E:** Comparison of performance metrics with and without 1p stimulation for Vglut1-Cre;Ai203 mice. Hit (blue) and False Alarm (brown) rates significantly increased (hit rate: p=0.019, false alarm rate: p=0.0016, paired t-test, n=4 mice) with photostimulation of excitatory cells. Points are individual mice, average of four sessions per mouse. **F:** Comparison of performance metrics with and without 1p stimulation for Vglut1-Cre;Ai203 mice. Hit (blue) and False Alarm (brown) rates significantly increased (Hit rate: no light 57% ± 1%, light 58 ± 2% p=.5; False alarm rate: no light 20 ± 1%, light 24 ± 2%, p=.25; paired t-test; n=3 mice) with photostimulation of excitatory cells. Points are individual mice, average of four sessions per mouse. Data are presented as mean ± bootstrapped 68% confidence interval.

## Methods

### Mouse transgenesis

The Ai203 transgenic mouse line was generated similarly as previously described ^25, 27^. The targeting vector was constructed using gene synthesis and standard molecular cloning techniques. It contains the following components: FRT3 – 2X HS4 chicken beta globin insulators – TREtight promoter – LoxP – ORF- 3X stops – hGH polyA, PGK polyA – LoxP – ChroME-FLAG-Kv2.1-GSGlinker-GcaMP7s– WPRE-bGH polyA – 2X HS4 chicken beta globin insulators – CAG promoter – Lox2272 – ORF-3X stops – hGH polyA, TK polyA – Lox2272 – nls-mRuby3-IRES2-tTA2 – WPRE – bGH polyA – PhiC31 attB- PGK promoter – Hygro1-SD – FRT5. Targeting of the transgene cassette into the TIGRE locus was accomplished via Flp-recombinase mediated cassette exchange using circularized targeting vector and a CAG-FlpE vector (Open Biosystems Inc) as previously described ^25, 27^. Correctly targeted ES cells were identified using PCR and injected into blastocysts to obtain chimeras and subsequent germline transmission. Resulting mice were then crossed to C57BL/6J mice and maintained in C57BL/6J congenic background. Only mice heterozygous for both reporter and driver transgenes (Slc17a7-IRES2-Cre;Ai203) were used for experiments.

### Histology

Mice were anesthetized with ketamine and transcardially perfused with phosphate-buffered saline (PBS) and then 4% paraformaldehyde (PFA). Brains were left in PFA overnight, then transferred to 30% sucrose for 1-2 days, then cut into 40 μm-thick coronal sections using a freezing microtome and stored in PBS until use.

For NeuN immunostaining, sections were blocked for 1 hour at 4 degrees on a rocker in PBS containing 3% normal goat serum (NGS), 0.6% Triton X-100, 0.2% Tween-20, and 3% bovine serum albumin (BSA). For primary staining, sections were incubated overnight at 4 degrees on a rocker in Rabbit anti-NeuN (Abcam) diluted 1:10,000 in blocking solution. The next day, sections were washed once in PBS with 0.25% Triton X-100 (PBS-T). For secondary staining, sections were incubated for one hour on the rocker at room temperature in blocking solution with 1:1000 Goat anti-Rabbit IgG H&L Alexa Fluor 405 (Thermo Fisher Scientific). Then sections were washed once in PBS-T. Sections were mounted and coverslipped with Vectashield (Vector Laboratories Inc).

Images were collected on an Olympus FV3000 (Olympus Corporation) confocal laser scanning microscope. For cell counting, to obtain adequate z-sectioning, fields of view tiled at 212 × 212 μm (800 × 800 pixels) each were taken spanning an area of V1, stitched in ImageJ with a stitching plugin ^79^. Then images were rotated so that the pia was horizontal and cropped so that the counting area was a rectangular column of cortex containing all layers, to evenly count all layers. Cells were counted in ImageJ using built-in tools.

For comparison to in utero electroporated st-ChroME-mRuby3, in utero electroporation was performed as described previously ^69^.

### Two-photon holographic microscopy

Slice electrophysiology and *in vivo* experiments were performed using two setups capable of 3D scanless holographic optogenetics with temporal focusing (3D-SHOT), as described previously ^41, 57, 69^. Both setups were custom-built on the Movable Objective Microscope (MOM; Sutter Instrument Co.) platform, with three combined optical paths: a 3D two-photon (2p) photostimulation path, a fast resonant-galvo raster scanning 2p imaging path, and a widefield one-photon (1p) epifluorescence/IR-transmitted imaging path, merged by a polarizing beamsplitter before the microscope tube lens and objective. In both setups, temporal focusing of the photostimulation beam from the femtosecond fiber laser was achieved with a blazed holographic diffraction grating (33010FL01-520R Newport Corporation). The beam was relayed through a rotating diffuser to randomize the phase pattern and expand the temporally focused beam to cover the area of the high-refresh-rate spatial light modulator (SLM; HSP1920-1064-HSP8-HB, 1920 × 1152 pixels, Meadowlark Optics). Holographic phase masks were calculated using the Gerchberg-Saxton algorithm and displayed on the SLM to generate multiple temporally-focused spots in 2D or 3D positions of interest ^80^.

The photostimulation path was then relayed into the imaging path with a polarizing beamsplitter placed immediately prior to the tube lens. As described in Mardinly et al., 2018, to limit imaging artifacts introduced by the photostimulation laser, the photostimulation laser was synchronized to the scan phase of the resonance galvos using an Arduino Mega (Arduino), gated to be only on the edges of every line scan.

The setup for *in vitro* characterization of Ai203 and 2P synaptic connectivity mapping used a Monaco 1035-80-60 737 (1040nm, 1MHz, 300fs, Coherent Inc.) for photostimulation and a Mai Tai Ti:sapphire laser (Spectra Physics Inc.) for 2p calcium imaging. For *in vivo* characterization of Ai203 mice, a similar setup was used except for a Monaco 1035-80-40 (1035nm, 2MHz, 276 fs, Coherent Inc.) was used for photostimulation, and a Chameleon Ultra Ti:sapphire laser (Coherent Inc.) was used for 2p calcium imaging.

### *In vitro* electrophysiology

*In vitro* slice recordings were performed on 300μm-thick coronal slices from both male and female mice aged P12-P30 (calcium sensor characterization) or P21-P42 (opsin characterization, and synaptic connectivity mapping experiments). Slice preparation was done as previously described ^57^. Whole-cell patch-clamp protocols were performed in inline heating-controlled (33°C) standard ACSF bath solution (in mM: NaCl 119, NaHCO_3_ 26, Glucose 20, KCl 2.5, CaCl 2.5, MgSO_4_ 1.3, NaH_2_PO_4_ 1.3). Patch pipette (4-7 MOhm) were pulled from borosilicate glass filaments (Sutter Instrument Co.) and filled with potassium (K)- gluconate solution (GCamp7s characterization; in mM: 110 K-gluconate, 10 HEPES, 1 EGTA, 20 KCl, 2 MgCl_2_, 2 Na-ATP, 0.25 Na-GTP, 10 Phosphocreatine, 295 mOsm, pH=7.45) or Cesium (Cs2+)-based internal solution (opsin characterization, and synaptic connectivity mapping experiments; in mM: 135 CeMeSO_4_, 3 NaCl, 10 HEPES, 0.3 EGTA, 4 Mg-ATP, 0.3 Na-GTP, 1 QX-314, 5 TEA-Cl, 295 mOsm, pH=7.45) also containing 50 μM Alexa Fluor hydrazide 488 or 594 dye (ThermoFisher Scientific). Data was recorded at 20 kHz using 700b Multiclamp Axon Amplifier (Molecular Devices). For loose-patch recordings the pipettes were filled with standard ACSF. The headstage with the electrode holder (G23 Instruments) was controlled by Motorized Micromanipulator (MP285A; Sutter Instrument Co.). All data was acquired and analyzed with custom code written in Matlab using the National Instruments Data Acquisition (DAQ) Toolbox (NI). Membrane (Rm) and series (Rs) resistance were monitored before and throughout experiments to ensure quality of acquired data. Cells were excluded if initial access resistance was over 30 MOhm or if access resistance changed more than 30% during the experiment, cells were excluded.

### In vitro characterization of intrinsic properties

Resting membrane potential was measured using K-gluconate internal solution and determined as the median membrane potential of 5 seconds of baseline measurement immediately following establishment of whole-cell configuration. Action potential threshold was analyzed using code (in MATLAB) adapted from Spike_threshold_PS function developed by the Rune Berg lab. This function determines threshold as the point of maximum positive slope in a phase plot of membrane potential and its first derivative dV/dt^81^.

Miniature EPSCs (mEPSC) were recorded at -70mV holding potential in bath solution containing tetrodotoxin (TTX: 0.5 μM) using Cs-based internal solution. A stable baseline was acquired for 3 minutes, followed by additional 5 minutes of recording. Postsynaptic events were detected and quantified across time bins using Easy Electrophysiology v.2.4.0 (Easy Electrophysiology Ltd), then statistical analyses were done using in-house code in MATLAB.

### *In vitro* characterization of opsin response characteristics

For 1p characterization, setup and methods were identical to those described for 1p characterization in Sridharan et al 2021. For 2p characterization, cells positive for st-ChroME-GCaMP7s were identified through fluorescence from widefield one-photon illumination using a Spectra X light engine (Lumencor). For both measurements of 2P-photoactivated peak currents (whole-cell voltage clamp) and spiking (loose-patch), the width of photostimulation was 5 ms. To measure peak 2P evoked photocurrents, a single holographic spot was aligned to a cell patched in whole-cell configuration and held in voltage clamp (-70mV) and stimulated across a series of laser powers (4.5, 7, 9, 13.5, 22.5, 45 mW) at 2 Hz. To measure spike probability, a cell in loose-patch configuration was stimulated with a single spot across laser powers (4.5, 7, 9, 11, 13.5, 22.5 mW) at different frequencies (5, 20, 40, 60 Hz). Stimulation conditions were randomized across an experiment.

The effective spatial resolution of photostimulation was measured in st-ChroME-GCaMP7s+ cells via loose-patch. We then measured spiking in response to photostimulation in the space surrounding the target cell by creating a voxelized three-dimensional volume spanning 97.5 × 97.5 × 50 μm in lateral and axial dimensions and at 6.5 × 6.5 × 12.5 μm per voxel. Spike probability was measured per stimulation of a voxel, then mapped according to the spatial location where the values were obtained. Lateral (x, y) and axial (z) physiological point-spread functions (PPSF) were quantified along grid planes where the highest spike probabilities per stimulated voxel were obtained. The spatial resolution at both lateral and axial dimensions were quantified as the full-width half-maximum (FWHM) of Gaussian profiles fitted to the PPSFs.

### In vitro imaging

For GCaMP characterization (Fig. 1), GCaMP7s positive cells were identified in 1P or 2P illumination and patched. After establishing stable access, cell parameters (Rm, Rs) were logged and 5 ms duration step current pulses of increasing amplitude were used to determine neuron rheobase. A train of 1, 2, 3, 5, 10, 15, 20 or 30 action potentials were triggered at 30 Hz, using 5 ms duration current pulses 25 pA higher than rheobase while simultaneously imaging calcium fluorescence at a 30 Hz frame rate. Similar imaging laser power was used (35 mW) for all cells. Imaging analysis is described with *in vivo* analysis.

Crosstalk characterization was performed as previously described ^69^, measured across FOV sizes (980 680 527μm), imaging frequencies (6, 10, 30 Hz), and powers (10, 15, 25, 35, 45 mW). Imaging frequencies were obtained by scanning the FOV at 30Hz while using the Pockels cell (Conoptics) to gate the scanning laser every n-th frame corresponding to frequency of interest. Cells were kept at resting potential throughout the protocol. Each trial consisted of a baseline period before two seconds of a scanning period.

Only the cells with stable resting potential (<4 mV change) over the protocol were accepted for data analysis. Data obtained were baseline-subtracted, then median-filtered to remove action potentials.

### *In vitro* synaptic connectivity mapping

Cells negative for st-ChroME-GCaMP7s (absence of fluorescence under 1P-illumination) were patched in whole-cell configuration and held at -70 mV. Preliminary assessment of photostimulation evoked postsynaptic responses and latency were done through widefield 1P stimulation at saturating power. To map synaptic inputs to the patched cell, the surrounding volume of 162.5 ×162.5 μm in lateral (x, y) and 100 μm in axial (z) dimensions was probed through holographic photostimulation of a precomputed three-dimensional grid of holographic spots. Each point (voxel) of photostimulation was spaced 6.5 × 6.5 × 25 (x, y, z) μm apart and were targeted randomly at 30 or 40 Hz with 5 ms laser pulses across 20 trials per laser power tested. Postsynaptic EPSCs within 25 ms time windows from the start of photostimulation at each voxel were collected and sorted. Synaptic connectivity maps were generated by plotting the peak of mean EPSCs according to the spatial location of the corresponding voxels.

### Viral injection and surgical procedures

For comparison with Ai203 reporter mice, double transgenic mice expressing CaMKII-tTA;tetO- GCaMP6s ^28^ were crossed with transgenic SST-Cre ^82^ or PV-Cre ^83^ mice to create triple-transgenic mice expressing GCaMP6s in excitatory (tTA-expressing) neurons. Mice were intracranially injected with PhP.eB-TRE-ChroME-P2A-H2B-mRuby3 ^84^, as previously described ^41^. Notably, as the tetracycline response element (TRE) viral promoter drives expression in cells expressing tetracycline trans-activator (tTA) ^85^, viral transduction was coincident with excitatory neurons transgenically expressing GCaMP6s, and not dependent on the particular Cre line used (PV-Cre or SST-Cre).

Mice 6 weeks or older were anesthetized with 2% isoflurane and administered 2 mg/kg of dexamethasone (to reduce cerebral edema) and 0.5 mg/kg of buprenorphine (as an analgesic). Mice were then secured in a stereotaxic frame (Kopf) and kept warm with a heating pad. A small incision was made to expose the skull and 2 burr holes were placed over left V1 (3.4 mm lateral, 1 mm anterior of lambda) approximately 1 mm apart. 300 nL of virus was slowly injected between 200-300 µm below the dura, at a rate of 50 nL/s using a microinjector (Micro4; World Precision Instruments) and a glass pipette (Drummond Scientific). The glass pipette was kept in place for 5-10 minutes after injection to ensure adequate spread and prevent backflow of virus during retraction of the pipette. Headplating and cranial windowing (below) were performed immediately after viral injection.

Mice were headplated with custom-made titanium headplates. To ensure secure placement, the skull was lightly etched with a scalpel and the headplate was initially attached over V1 with Vetbond (3M). Metabond (C&B) and OrthoJet (Lang Dental) were used to permanently attach and secure the headplate. A craniotomy was made over the injection sites and V1 using a 3.5 mm biopsy punch. Once the skull was removed, bleeding was controlled with cold phosphate-buffered saline and Gelfoam (Pfizer Inc.). A cranial window was constructed by gluing together two 3 mm diameter circular coverslips to the bottom of a 5 mm diameter circular coverslip, then placed onto the craniotomy, and secured into place with Metabond (C&B). Mice were allowed to recover in a heated recovery cage.

### *In vivo* imaging and photostimulation

For 2-photon calcium imaging *in vivo*, mice were head-fixed and allowed to run on a freely moving circular treadmill under a 20X magnification (1.0 NA) water-immersion objective (Olympus Corporation) and imaged with a Sutter MOM 2p resonance scanning microscope controlled with MATLAB-based ScanImage software (Vidrio Technologies) (see also microscope design, above). Fast, 3 or 4 z-plane volumetric imaging (3 planes: 6.36 Hz, 4 planes: 4.77 Hz, field-of-view: 800 µm x 800 µm) was accomplished using an electrically-tunable lens (Optotune) placed in the light path prior to the resonance scanners. A 20 ms delay was added to the Y-galvo flyback time to provide adequate settle-time for the ETL and ensure a non-distorted field-of-view (FOV). Z-planes were placed 20 or 30 µm apart for 4 or 3 plane recordings, respectively. The imaging laser (Chameleon; Coherent Inc.) was tuned to 920 nm and restricted to <75 mW average power to limit scanning-induced crosstalk.

ChroME-expressing cells were identified by nuclear-localized mRuby3 fluorescence imaged at 1040 nm and using a custom 2d-peak detection algorithm in MATLAB. Detected cell locations were used as targets for holographic photostimulation. Custom written MATLAB code was used to synchronize the experiment via NI data acquisition boards.

Visual stimuli were presented on a 2048 x 1536 Retina iPad LCD display (Adafruit Industries) placed 10 cm from the mouse. The monitor backlight was synchronized with the galvos such that it came on only during the turnaround time, so that light from the monitor did not contaminate 2p imaging. Visual stimuli were created and presented with custom MATLAB code and Psychophysics ToolBox. Drifting gratings (50 visual degrees, 1 Hz, .08 cycles per degree, 100% contrast) of different orientations were randomly presented for 1 second each trial and interleaved with a grey-screen (“blank”) condition. Neurons with significantly different responses to visual stimuli (p<0.05, 1-way ANOVA) were considered as visually responsive. FOVs with fewer than 20% visually responsive cells were considered unlikely to be in V1 and excluded from further analysis. Preferred orientation was calculated as the max response from trial-averaged orientation tuning curves; orthogonal orientation was determined as the response 90 degrees from preferred orientation. Orientation selectivity index (OSI) was then calculated as:

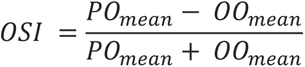

Where PO is the mean response to preferred orientation and OO is the mean response to orthogonal orientation. Orientation tuning curves were min subtracted prior to OSI calculation to ensure values ranged from 0-1.

To construct contrast response curves, **full-screen Gaussian contrast modulated noise stimuli (CITE NEEDED)** were presented using custom written MATLAB scripts and Psychophysics Toolbox. Contrast response functions were fit with a modified Naka-Rushton function^86^, defined as:

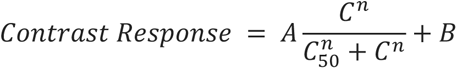

Where A is the maximum firing rate of the neuron, B is the baseline firing rate, C is the contrast shown, and C_50_ is the contrast corresponding to 50% maximum response, and n is an exponent controlling the slope. Contrast response curves were fit by least-squares regression using SciPy’s curve_fit function in Python. B was set constant to 0, corresponding to the low baseline firing rates of pyramidal cells *in vivo*. n was constrained between 0 and 4.

### Power and Spike Calibration

All data was processed online via a real-time implementation of CaImAn OnACID^87^ coined ‘live2p’ (www.github.com/willyh101/live2p). Briefly, the autodetected red nuclei (or, in some cases, manually selected green cells) used as targets for photostimulation were used as seed locations for fluorescent sources. Images were streamed from ScanImage Tiffs into multithreaded processing queues for each plane, allowing for simultaneous processing of multiple imaging z-planes in separate threads. Motion correction and signal extraction was performed during the experiment only on cells used for seeding to improve online performance.

Throughout all photostimulation experiments, all multi-target holograms are corrected for diffraction efficiency and overall efficiency, as measured by calibration of 3D-SHOT^57^.This ensures each target receives the intended power regardless of where its located, or what other targets are present.

A subset of detected cells were power calibrated. Groups of 5 targets per hologram were driven with five 5ms pulses at 30Hz. Targets were arranged in holograms to maximize the distance between any two simultaneously shot targets. A ramp of powers from 0 up to 85mW (4 to 12 steps total) was used, with 5 to 10 repetitions per power step. Afterward, mean denoised, neuropil-decontaminated fluorescence, as measured by OnAcid, one second after stimulation was taken. The highest and lowest fluorescence trial for each power step were excluded to handle outliers. The data was fit with a ‘hill function’, below.

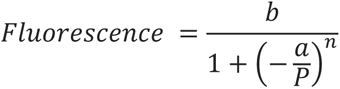

Where b represents the fluorescence at saturation, a is the power that produces 50% of the saturating response, P is the optogenetic stimulation power, and n can be either 4, 6, or 8 and controls the slope of the function. The ‘target power’ is defined as the power needed to reach 80% of the saturating fluorescence (b). Cells were considered not fit and excluded from further analysis if R^2^ < 0.25, or the ‘target power’ was below 10mW or above 1.5x the max power tested.

Cells that were fit by power calibration and were visually responsive were spike calibrated using a similar design. As before, cells were grouped into holograms containing 5 spread out targets, with each stimulated at 10 – 20% more than their fit target power. Each cell was stimulated for 1s with 1 to 20 pulses. The observed fluorescence was fit as a function of spike number by a quadratic function with the 0-spike intercept fixed at 0 fluorescence. Cells with an R^2^<0.15 were considered not fit and excluded.

Next, we presented visual stimuli, using the protocol described above, in 8 directions and measured response to each. Stimuli were 1 s long. Neural responses were measured as the mean responses in the 1 second during visual stimulation. We used the fit model from spikes to calcium to estimate the number of optogenetic pulses it would take to match the observed mean fluorescence from the visual stimulus.

To generate a multi-cell pattern, a one second stimulation period was divided into 50 holograms encoding 0 to 10 targets each and triggered at 50Hz. A single hologram can activate at least 50 cells simultaneously (Fig. 2I-J), and with a 300 Hz SLM we can stimulate and switch holograms at 125Hz, allowing a wide variety of large-scale population vectors to be written^41, 77^. A custom built gradient descent discrete optimizer was designed to calculate the precise pattern of which cells are stimulated in which hologram. After assigning the desired number of spikes per cell, the cost function of this optimizer attempted to maximize each cell’s ISI, while keeping the number of targets per hologram relatively low.

Two target holograms were prevented, as there it is difficult to deliver the correct power to both targets. One target holograms often deliver more power than multi-target holograms, and therefore were penalized but not prevented. Each hologram was triggered 5ms before a 5ms optogenetic pulse, each cell received 10-20% more power than the target power. Ensembles contained 19 to 43 cells with up to 168 spikes added in total.

### Calcium imaging post-processing and analysis

For figures 1-2, calcium imaging data was extracted from raw TIFF files, motion corrected, and source-extracted using Suite2p ^88^. Putative sources were manually curated based on morphology and fluorescence statistics. Neuropil traces for each source were generated by Suite2p, multiplied by a correction coefficient (0.7), and subtracted from source fluorescence traces. Calcium traces were minimum subtracted, and ΔF/F was calculated as:

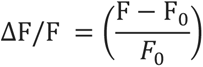

Where F is a cell’s fluorescence response and F_0_ is baseline fluorescence. For *in vitro* characterization in Figure 1, F_0_ was set to be the mean fluorescence one second pre-stimulus. For *in vivo* characterization in Figure 2, F_0_ was calculated by a 1-minute rolling 20% quantile. Individual trialwise responses were baseline subtracted using the mean pre-stimulus (holographic or visual) activity.

Holographic targets were matched to sources identified by Suite2p by the source satisfying the *argmin* Euclidean distance for each target. Targets lacking a source within 15 µm were excluded from further analysis. In the rare cases where multiple targets matched to the same source, only the closest target was used.

For *in vitro* characterization in Figure 1, zero stimulation trials were simulated by taking fluorescence from the inter-trial interval of one spike trials, six seconds after the last spike stimulation. Mean calcium traces were calculated by taking the mean calcium trace across all trials for a given spike number for a given cell. Peak ΔF/F was the peak of the mean calcium trace, with the peak ΔF/F for 0 spike trials subtracted to account for noise.

Cells were considered to be “stimmable” if their mean response to holographic stimulation was significantly different (p<0.05, one-tailed rank-sum) from baseline.

For Figures 3-4, Because suite2p uses a different method than CaImAn for cell identification and analysis, for consistency with online measurements taken during calibration, to analyze the power and spike calibration data, and the writing in of population vectors after power and spike calibration, we used live2p/CaImAn OnACID run offline on the entire imaging session at once, seeded to our targets. This motion corrected and source-extracted the data, and used the z-scored neuropil-corrected and denoised calcium signal for analysis. We also analyzed the data using suite2p, as and obtained similar results. In fact, we found that with suite2p, a higher % of matched targets were fit in the power balancing step, 80.5% (803/997 detected cells) were fit resulting in a similar mean target power (66.3 ± 0.9 mW). Thus, we believe that most of the cells that were not fit represent low signal to noise cells or ROIs that were falsely identified as cells.

### Visual contrast detection task

#### Apparatus

The visual behavioral apparatus for the contrast detection task was controlled using an Arduino DUE synchronized with Raspberry Pi 3+/4+, which interfaced with a custom-written code in Python, Arduino and Java. Mouse licking was detected by changes in the capacitive load using a capacitive touch sensor (Adafruit, AT42QT1070) forming a circuit between the 0.05-inch diameter steel lickport and the mouse’s tongue. Water was delivered by gravity though the lick port using a 2- way normally closed isolation valve control (Neptune Research Inc.). Visual stimuli were generated using a custom-written code in Python and presented using a gamma corrected LCD monitor (Podofo, 15.7 x 8.9 cm, 1024x600 pixels, 60 Hz refresh rate) located 12 cm from the right eye. The entire apparatus was mounted on an 8.0 x 8.0 x 0.5 inch (MB8; Thorlabs Inc.) aluminum breadboard and enclosed in a light isolation box (80/20).

#### Behavioral Training

Training began after headplate and cranial window implantation surgery, recovery (7 days) and water restriction (5-8 days). Prior to training, mice were habituated to head fixation in the tube apparatus with the monitor on. The habituation sessions were performed for 2 days lasting ∼30 minutes each. The training schedule was divided into four main stages. During the first stage of behavioral training, head fixed mice were classically conditioned to lick in response to the presence of a visual stimulus consisting of a square patch of a drifting sinusoidal grating (2Hz, 0.08 cycles/degree, 600 ms, 100% contrast, 34 visual degrees, black background luminescence). On every trial, the visual stimulus was paired with a water reward delivered at the beginning of the response window. No catch trials occurred in this stage. Once mice reliably licked correctly within the response window in more than 80% of the trials, they moved onto the operant conditioning phase of training.

During the second stage of training, mice were trained to respond in the presence of the visual stimulus. Catch trials were introduced as 25% of trials. The mouse must lick in response to the visual stimulus within the reward window to receive a water reward and withhold licking otherwise to avoid time out punishments. Mice started this paradigm with the same visual stimuli features as stage 1(100% contrast, 34 VD, black background luminescence). Once mice reached performance criteria (>90% hit rate and <30% false alarm rate) for two consecutive days, mice were transitioned to stage three.

In stage three, background luminescence was ramped up until it reached mean luminance, then stimulus size was ramped down until it reached 18 VD. Ramping occurred within and across sessions as long as performance criteria were maintained. Once ramping was complete, mice had to maintain performance at criteria for one day to progress to the next stage.

In stage four, mice had to detect stimuli of varying contrasts. For strain comparisons (Fig. S6B-C), 10 different contrasts (not including 0% catch trials) were used and catch trials were 7.5% of all trials. For optogenetic testing (Fig. S6D-F), six different contrasts were used and catch trials were 25% of all trials.

In the fourth stage of training, mice started to detect visual stimuli with six different contrasts, customized for each mouse, to provide a reliable behavioral threshold. Stimuli of different contrasts were randomly interleaved within blocks. Catch trials represented 12% of the trials.

#### Task Design

The contrast detection task consisted of uncued GO or CATCH trials that occurred after random intertrial intervals (3-8 seconds). A no lick period was enforced in the two seconds preceding the trial, licking during this period caused a time-out punishment (5-9 seconds). In GO trials a visual stimulus of variable contrast was presented, if the mouse responded within the 500 ms response window, it received a reward. CATCH trials had identical timing, but no stimulus was presented and if the mouse licked within the response window it received a time out punishment. Time outs were 5-9 seconds. The 500 ms response window immediately followed the 600 ms stimulus presentation. Contrasts were presented in randomly shuffled blocks wherein each contrast must occur three times.

#### Strain information

Wildtype mice were C57BL/6J mice. Camk2a-tTA;tetO-GCaMP6s mice were double transgenic heterozygotes.

### One-photon optogenetic stimulation

One-photon (1p) photostimulation during contrast detection task was delivered using an optical fiber (400µm diameter, 0.39 nA; Thorlabs Inc.) coupled to a 470-nm LED (M470F3, Thorlabs) driven by an LED driver (LEDD1B; Thorlabs Inc.) and positioned using a micromanipulator over the location of V1 at the posterior of the window. Light power was calibrated at the end of the fiber using a power meter (PM160T, Thorlabs). Stimulation power was calibrated to each mouse and ranged from 0.05-0.4 mW out of the fiber. For control wildtype mice, 0.5 mW was used for all mice. To prevent photostimulation from interfering with behavioral performance, we attached a custom designed light-blocking cone to the mouse head and illuminated blue LED masking lights during the entire session. To further reduce any extraneous cues, we covered the left eye with a plastic eye patch. Light was delivered from the onset of the stimulus to the end of the response window on 33% of trials.

## Supplementary note

In the TIGRE2.0 system, two separate transgene-expressing units are co-inserted into the TIGRE locus ^25^. Both are Lox-Stop-Lox (LSL) controlled, but the first is TRE driven and the second is CAG driven. In Ai203, the st-ChroME-GCaMP7s is TRE-driven, while the second CAG-driven unit contains an nls- mRuby3 and the tTA2 that drives the first unit (full gene: TITL-st-ChroME-GCaMP7s-ICL-nls-mRuby3- IRES2-tTA2; Fig. 1A).

Certain combinations of Cre-driver and TIGRE2.0 reporter lines drive expression of the TRE-driven protein in only a subset of Cre+ cells, most notably in crosses containing Pvalb-Cre ^25^. In other TIGRE2.0 reporter lines with one TRE-driven fluorophore and one CAG-driven fluorophore, the CAG-driven fluorophore has been expressed in almost all the target population even when the TRE-driven fluorophore is sparsely expressed relative to the expected Cre+ population. This is thought to be due primarily to promoter silencing of the TRE promoter, not due to a lack of tTA2. Consistent with this, the CAG-driven nls-mRuby3 in Ai203 was densely expressed (74 ± 4% of all anti-NeuN labelled neurons), while the TRE-driven st-ChroME- GCaMP7s was sparser.

This sparsity has not been noted in excitatory cortical cells before. Unlike previously published TIGRE2.0 lines, Ai203 has an IRES2-tTA2 and uses the TREtight rather than the TRE2 promoter. The IRES2 will likely decrease expression of the tTA2, which may reduce overall expression levels, but could lead to greater overall health. TREtight is more prone to silencing, so this may lead to the lower expression seen in Ai203 compared to other TIGRE2.0 lines.

